# SARS-CoV-2 Omicron spike mediated immune escape and tropism shift

**DOI:** 10.1101/2021.12.17.473248

**Authors:** Bo Meng, Isabella A.T.M Ferreira, Adam Abdullahi, Niluka Goonawardane, Akatsuki Saito, Izumi Kimura, Daichi Yamasoba, Pehuén Perera Gerba, Saman Fatihi, Surabhi Rathore, Samantha K Zepeda, Guido Papa, Steven A. Kemp, Terumasa Ikeda, Mako Toyoda, Toong Seng Tan, Jin Kuramochi, Shigeki Mitsunaga, Takamasa Ueno, Kotaro Shirakawa, Akifumi Takaori-Kondo, Teresa Brevini, Donna L. Mallery, Oscar J. Charles, CITIID-NIHR BioResource COVID-19 Collaboration, The Genotype to Phenotype Japan (G2P-Japan) Consortium, Ecuador-COVID19 Consortium, John E Bowen, Anshu Joshi, Alexandra C. Walls, Laurelle Jackson, Sandile Cele, Darren Martin, Kenneth G.C. Smith, John Bradley, John A. G. Briggs, Jinwook Choi, Elo Madissoon, Kerstin Meyer, Petra Mlcochova, Lourdes Ceron-Gutierrez, Rainer Doffinger, Sarah Teichmann, Matteo Pizzuto, Anna de Marco, Davide Corti, Alex Sigal, Leo James, David Veesler, Myra Hosmillo, Joo Hyeon Lee, Fotios Sampaziotis, Ian G Goodfellow, Nicholas J. Matheson, Lipi Thukral, Kei Sato, Ravindra K. Gupta

## Abstract

The SARS-CoV-2 Omicron BA.1 variant emerged in late 2021 and is characterised by multiple spike mutations across all spike domains. Here we show that Omicron BA.1 has higher affinity for ACE2 compared to Delta, and confers very significant evasion of therapeutic monoclonal and vaccine-elicited polyclonal neutralising antibodies after two doses. mRNA vaccination as a third vaccine dose rescues and broadens neutralisation. Importantly, antiviral drugs remdesevir and molnupiravir retain efficacy against Omicron BA.1. We found that in human nasal epithelial 3D cultures replication was similar for both Omicron and Delta. However, in lower airway organoids, Calu-3 lung cells and gut adenocarcinoma cell lines live Omicron virus demonstrated significantly lower replication in comparison to Delta. We noted that despite presence of mutations predicted to favour spike S1/S2 cleavage, the spike protein is less efficiently cleaved in live Omicron virions compared to Delta virions. We mapped the replication differences between the variants to entry efficiency using spike pseudotyped virus (PV) entry assays. The defect for Omicron PV in specific cell types correlated with higher cellular RNA expression of TMPRSS2, and accordingly knock down of TMPRSS2 impacted Delta entry to a greater extent as compared to Omicron. Furthermore, drug inhibitors targeting specific entry pathways demonstrated that the Omicron spike inefficiently utilises the cellular protease TMPRSS2 that mediates cell entry via plasma membrane fusion. Instead, we demonstrate that Omicron spike has greater dependency on cell entry via the endocytic pathway requiring the activity of endosomal cathepsins to cleave spike. Consistent with suboptimal S1/S2 cleavage and inability to utilise TMPRSS2, syncytium formation by the Omicron spike was dramatically impaired compared to the Delta spike. Overall, Omicron appears to have gained significant evasion from neutralising antibodies whilst maintaining sensitivity to antiviral drugs targeting the polymerase. Omicron has shifted cellular tropism away from TMPRSS2 expressing cells that are enriched in cells found in the lower respiratory and GI tracts, with implications for altered pathogenesis.

## Introduction

The Omicron variant, first detected in South Africa, carries over 30 mutations in its spike protein and has now spread internationally at ferocious pace^1^. More than 20 substitutions exist on the N-terminal domain (NTD) and receptor binding domain (RBD), which are the main targets for neutralising antibodies. The Omicron variant harbours six unique mutations in S2 (N764K, D796Y, N856K, Q954H, N969K, L981F) that were not previously detected in other variants of concern (VoC). Alpha and Delta variant spike proteins were previously shown to confer more efficient cell-cell fusion kinetics compared to Wuhan-1^2^ due to mutations in the furin cleavage site region that increase S1/S2 cleavage, and syncytia formation previously associated with pathogenesis^3^. Omicron has three mutations in this region (P681H, H655Y and N679K) and therefore was initially predicted to be highly infectious^4^ and pathogenic^5^. Indeed, the Omicron variant has been associated with very rapid increases in case numbers and recent data demonstrate significant re-infection and vaccine ‘breakthrough’, likely due to evasion of neutralising antibody responses^1,6^. Somewhat paradoxically however, recent findings suggest reduced severity in Omicron infections compared to Delta^7^.

Here we explore biological properties of Omicron, and focus on spike mediated evasion of neutralising antibodies, increased ACE2 binding affinity, as well as shift in tropism away from TMPRSS2 expressing cells and compromised ability to generate syncytia.

## Results

### Omicron spike binds ACE2 with enhanced affinity

The Omicron variant has 15 amino acid mutations in the RBD (Figure 1a-b and Supplementary Figure 1), three of which (K417N, T478K, and N501Y) have also been observed in previous VoCs. To understand the impact of these substitutions on receptor engagement, we determined the kinetics and affinity of monomeric human ACE2 binding to immobilized Omicron, Wuhan-Hu-1 and Alpha RBDs using biolayer interferometry (BLI). We observed that the Omicron RBD has a 2-2.5-fold enhanced binding affinity for ACE2 relative to the Wuhan-Hu-1 RBD (Table 1, Supplementary Figure 5), in line with recent surface plasmon resonance findings^8,9^.As previously observed for the Alpha RBD, which only harbors the N501Y mutation^10^, the modulation of binding is mediated by changes in ACE2 binding off rates. Although the K417N mutation is known to dampen ACE2 engagement^10,11^, a recently determined crystal structure of ACE2-bound Omicron RBD revealed that Q493R and Q498R introduce additional electrostatic interactions with ACE2 residues E35 and D38, respectively^11^, whereas S477N enables hydrogen-bonding with ACE2 S19. Collectively, these mutations strengthen ACE2 binding, relative to the ancestral isolate. We extended binding analysis to a cell based model using cells transfected with full-length spike, followed by ACE2 antibody titration; we observed significantly higher ACE2 binding for Omicron spike as compared to both Wuhan-1 and Delta variant spikes (Supplementary Figure 5). This enhanced binding could be a factor in the enhanced transmissibility of Omicron relative to previous variants.

**Figure 1.**
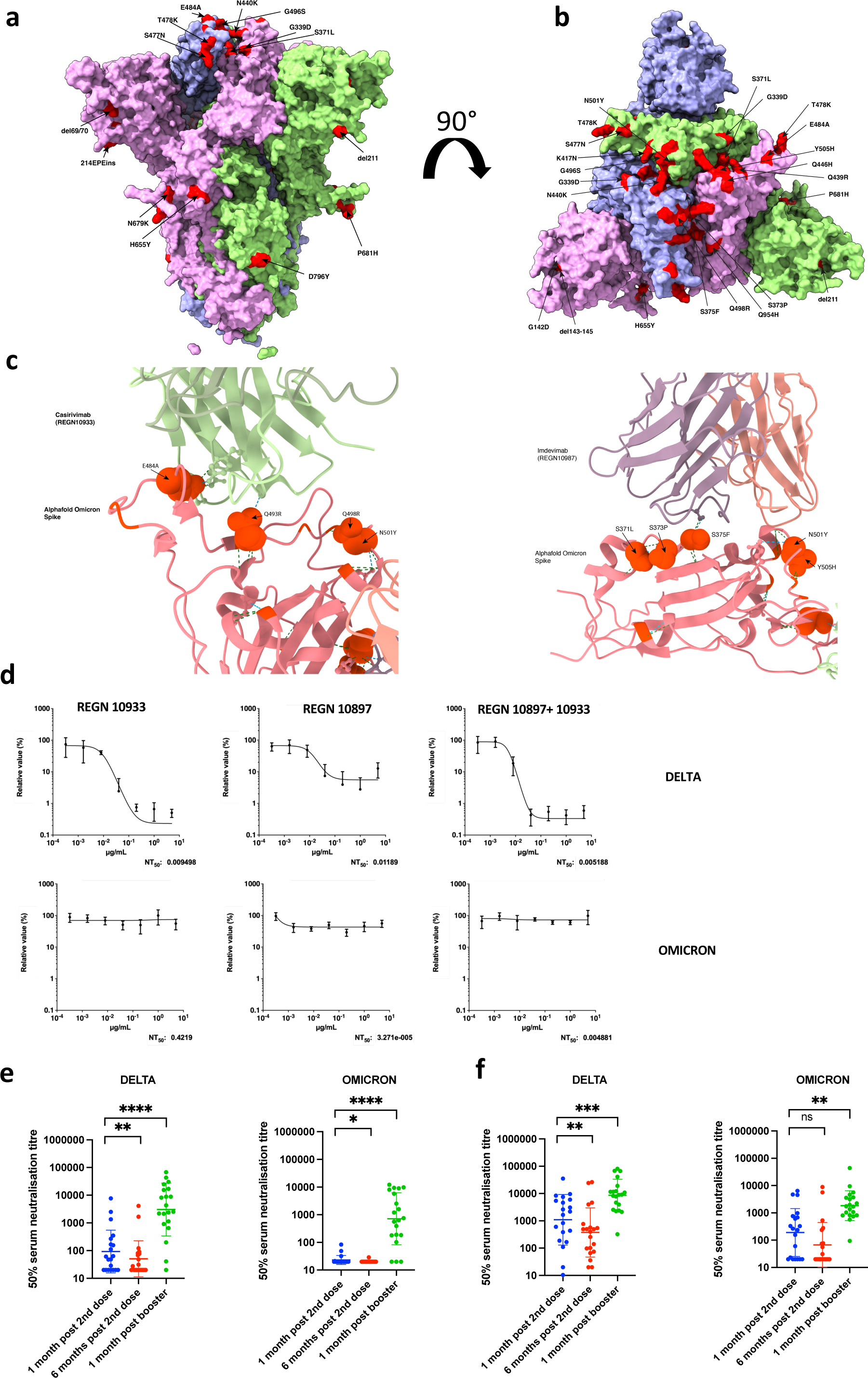
Sensitivity of SARS-CoV-2 Omicron to clinically approved monoclonal antibodies directed against spike and to vaccine elicited neutralizing antibodies. a. Side Surface representation of the Omicron spike protein. b. Top down surface representation of the Omicron spike. Spike homotrimer structures were created predicted *in silico* by the Alphafold2 software package. Individual mutations making up the Omicron spike are highlighted in red on each of the three homotrimers. c. Predicted interaction sites for REGN 10933 and 10897 monoclonal antibodies with Omicron spike RBD. Hydrogen bonds are indicated with dashed lines. Predicted contact points are shown as spherical representation. Omicron mutations labelled. d. titration of monoclonal antibodies REGN 10933 and 10897 and combination against replication competent Delta and Omicron viruses. NT: Neutralising titre. e.Neutralisation of spike pseudotyped virus by sera from vaccinated individuals over three time points following dose two (ChAdOx-1 or BNT162b2) and dose three (BNT162b2 only) e. n=20 ChAdOx-1 or f. n=20 BNT12b2. GMT (geometric mean titre) with s.d are presented. Data representative of two independent experiments each with two technical replicates. **p<0.01, *** p<0.001, ****p<0.0001 Wilcoxon matched-pairs signed rank test, ns not significant.

**Table 1.** Kinetic Analysis of human ACE2 binding to SARS-CoV-2 VOC by Biolayer Interferometry. Values reported represent the global fit to the data shown in Supplementary Figure 5.

|  | Wuhan | Alpha | Omicron |
| --- | --- | --- | --- |
| **KD (nM)** | 137 | 26 | 63 |
| **k<sub>on</sub> (M<sup>-1</sup> s<sup>-1</sup>)** | 1.2 x 10<sup>5</sup> | 1.2 x 10<sup>5</sup> | 1.1 x 10<sup>5</sup> |
| **k<sub>off</sub> (s<sup>-1</sup>)** | 1.6 x 10<sup>-2</sup> | 3.0 x 10<sup>-3</sup> | 7.0 x 10<sup>-3</sup> |

**Table 2:** Demographic characteristics of participants in vaccine elicited sera neutralisation study. Categorical variables (expressed as proportions and percentages) and continuous variables (expressed as medians with inter quartile ranges [IQR]) were compared using chi-squared test or Wilcoxon Mann Whitney tests, as applicable.

| Characteristic | Total | Pfizer (BNT162b2) | AstraZeneca (ChAdOx1) |
| --- | --- | --- | --- |
| Total number, n (%) | 40 | 20 (50) | 20 (50) |
| Age, median years (IQR) | 70.5 (62.5, 74.0) | 71.0 (55.8, 75.8) | 69.5 (63.0, 73.8) |
| ≥80, n (%) | 4 (10) | 4 (20) | 0 (0) |
| <80, n (%) | 36 (90) | 16 (80) | 20 (100) |
| Gender, n (%) |  |  |  |
| Male | 15 (37.5) | 8 (40) | 7 (35) |
| Female | 25 (62.5) | 12 (60) | 13 (65) |
| Previous COVID-19 |  |  |  |
| Yes | 5 (12.5) | 2 (10) | 3 (15) |
| No | 35 (87.5) | 18 (90) | 17 (85) |
| Type of booster |  |  |  |
| Moderna | 3 (7.5) | 0 | 3 (15) |
| Pfizer | 37 (92.5) | 20 (100) | 17 (85) |
| Duration between booster dose and sampling, median days (IQR) | 34 (32, 36) | 34.0 (32.0, 36.0) | 33.5 (30.3, 35.8) |

### Omicron escapes from a clinically approved combination monoclonal antibody therapy

Omicron was predicted to have broad resistance to neutralising antibodies based on mutations in class I-IV antigenic regions in the RBD^12^. Current standard of care antiviral treatment for moderate to severe COVID-19 includes use of the monoclonal antibody combination REGN 10933 and 10897. In the absence of clinical data for efficacy of these treatments against Omicron, we first modelled the interaction surface between each antibody and Omicron spike using published structures (Figure 1c). The E484A and Q493R changes were predicted to impact interactions with casivirimab and S375F with imdevimab. We next tested serial dilutions of component mAbs, both individually and in combination, against Delta and Omicron live viruses in tissue culture (Figure 1d). Whilst the Delta variant was effectively neutralised by casivirimab, imdevimab was partially effective, consistent with previous data^13^. The combination was highly potent against Delta. However, there was complete loss of neutralising activity against Omicron by either mAb alone or in combination (Figure 1d). Given these results, we next tested direct acting antivirals remdesivir and the active metabolite of molnupiravir against live virus. We observed similar antiviral activity against Delta and Omicron using serial titrations of both compounds (Supplementary Figure 2).

### Omicron spike protein confers broad escape from two, but not three, dose vaccination

A critical question is whether vaccine elicited antibodies are able to neutralise Omicron. We synthesised codon optimised spike expression plasmids for Omicron and Delta spike proteins and generated PV particles by co-transfecting the spike expression plasmids with a lentiviral gag-pol expressing plasmid and a lentiviral transduction plasmid encoding the luciferase gene^14,15^. We obtained longitudinal serum samples from 40 individuals vaccinated with either BNT162b2 or ChAdOx-1 vaccines and performed serum titrations before mixing sera with our reporter PV particles. Participants had median age of around 70 years and prospective serum samples were taken as follows: one month after dose two, six months after dose two, and one month after dose three (Supplementary table 2). We observed ≥ 10-fold loss of neutralisation against Omicron after the second dose compared to Delta (Figure 1e, f, Supplementary Figure 3). Indeed neutralisation of Omicron was not detectable for the majority of individuals who had received two doses of ChAdOx-1. We additionally also observed waning over time since second dose (Figure 1e,f, Supplementary Figure 3). Both groups were boosted with BNT162b2 as a third dose, allowing us to compare the response to this boosting dose. Very substantial increases (> 10 fold) in neutralisation against both Omicron and Delta were observed for all variants following third dose vaccination, suggesting increased breadth of responses as well as titre.

To confirm loss of neutralising activity against Omicron after the second dose, we next used a live virus experimental system to compare Delta and Omicron variants against sera taken four weeks after the second dose of BNT162b2, with similar results as obtained in the PV assay (Supplementary Figure 4a). These live viruses were also used to assess the neutralisation of the Omicron variant by sera derived from non-vaccinated individuals previously infected with the early Wuhan-1 virus, or the Delta variant. As expected, vaccine sera had significantly impaired activity against Omicron as compared to Delta (Supplementary Figure 4b). We also tested mRNA 1273 vaccine elicited sera which showed similar fold neutralisation loss to BNT162b2. Coronavac sera however showed little neutralisation against Delta and 0/9 participants had detectable neutralisation against Omicron. Interestingly sera from Delta infections appeared to have lower cross neutralisation as compared to those from the early pandemic period when Wuhan-1 D614G was dominant (Supplementary Figure 4b).

### Omicron S1/S2 cleavage is inefficient with impaired replication in certain cell types

We infected primary human nasal epithelial 3D cultures (hNEC) with an air liquid interface (Figure 2a,b). Infection with live SARS-CoV-2 Omicron or Delta was conducted at the apical surface with equal amounts of input virus and virus collected from the apical surface at 24 and 48 hours, with quantification of virus by E gene qPCR and TCID50. We observed similar replication kinetics for Omicron and Delta in the hNEC (Figure 2b). We next infected Calu-3 lung cells (known to express endogenous TMPRSS2) with live isolates, and observed significantly greater viral replication for Delta than Omicon (Figure 2c), manifesting as early as 24 hours post infection. We also infected Caco2, a colon cancer cell line (known to express high levels of endogenous TMPRSS2^16^), and, as for Calu-3, found greater Delta RNA and infectious virus (TCID50) in supernatants than was observed for Omicron. Finally we tested a cell line, HeLa, overexpressing ACE2 and TMPRSS2 with similar results as for Calu-3 and CaCo-2 cells.

**Figure 2:**
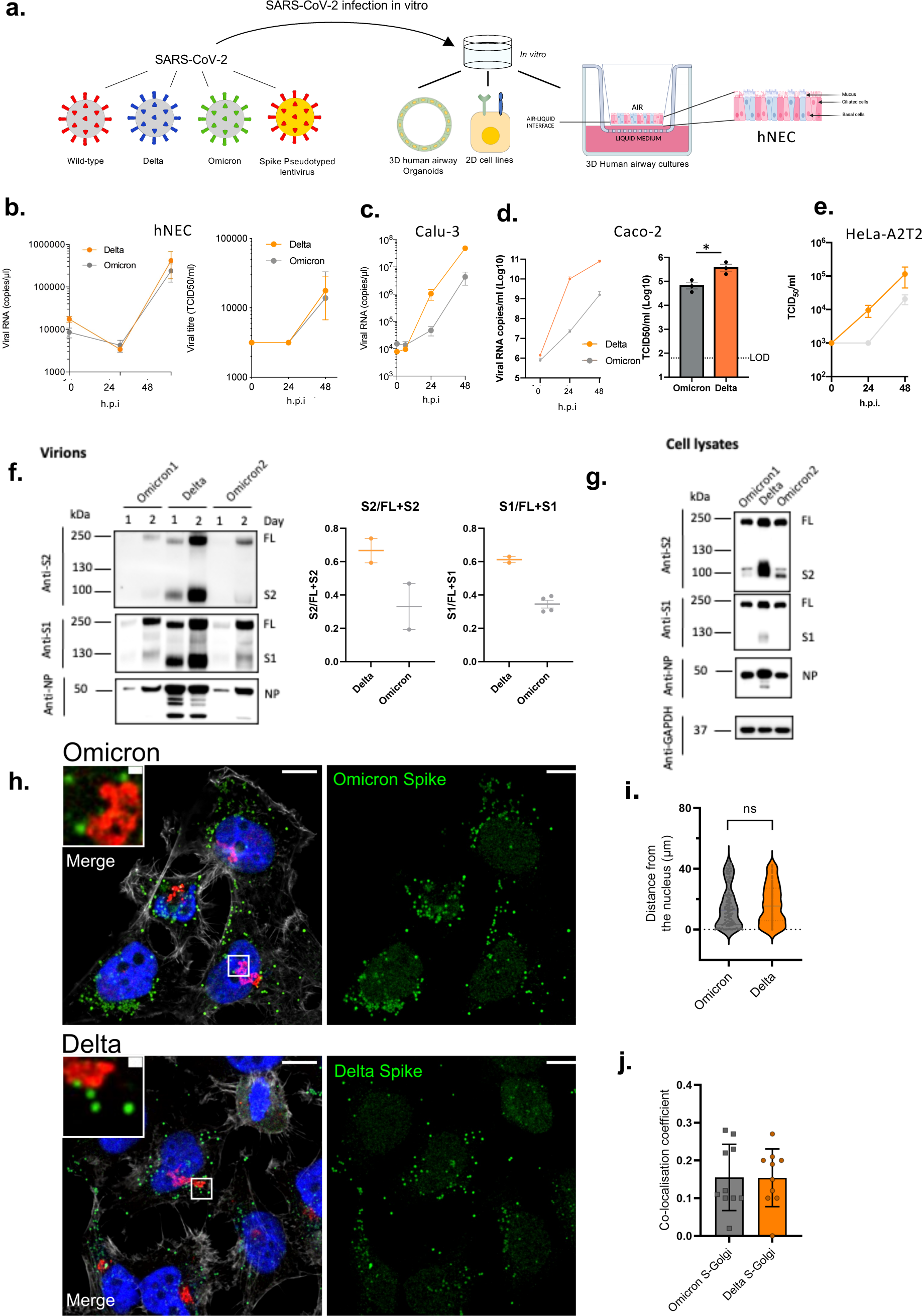
SARS-CoV-2 Omicron and Delta Variant live virus replication in 3D tissue culture systems and 2D cell lines. **a. overview of viruses and culture systems used hNEC (human nasal epithelial cultures). b-d**. Spreading infection by replication competent Omicron versus Delta variant over time in **b**. hNEC **c**. Calu-3 **d**. Caco-2 Npro **e**. HeLa-A2T2. Viral RNA and/or infectious virus in supernatant (TCID50) were measured **f-g**. Western blot of **f**. Two live Omicron and one Delta virus isolates probed with antibodies to S2, S1 and NP with quantification of S2 and S1 to total spike ratio. **g**. Vero E6 A2T2 producer cell lysates infected with live isolates probed with antibodies to S2, S1 and NP. Housekeeping gene GAPDH as loading control. Data are representative of two independent experiments. **h-j. Subcellular localisation of Spike in SARS-Cov-2 Delta vs Omicron infected cells**. Subcellular distribution of Omicron, Delta Spike proteins in Hela-ACE2 cells infected with live virus isolates. **h**. Cells on coverslips were infected for 24 h, fixed and stained with anti-Spike, anti-GM130-cis-Golgi, phalloidin 647 and DAPI, and imaged on a Leica TCS SP8 confocal microscope. **i**.The distance of spike proteins from nucleus at 24 hpi. **j**. Quantitation of Spike-Golgi colocalisation in infected cells. Values were calculated using Pearson’s correlation coefficient. Scale bars: 10 μm and 1 μm, respectively. * p<0.05; NS – not significant

We next sought to examine the possibility that the impaired replication in some cells may be related to differential spike incorporation and/or cleavage. For example Delta, known to have higher replication, is associated with a highly cleaved spike protein^2^. We tested purified virions from two independent Omicron isolate infections. We observed broadly similar amounts of Omicron and Delta spike incorporation into virions generated in VeroE6 ACE2/TMPRSS2 cells (Figure 2f). There was, however, reduced cleavage in whole Omicron virions compared to Delta as evidenced by the ratio of full length spike to S1; the Omicron spike was inefficiently cleaved in virions compared to Delta (Figure 2f). However, a more dramatically reduced Omicron spike cleavage relative to Delta was observed in cell lysates (Figure 2g).

Given lower intracellular S1/S2 cleavage for Omicron versus Delta spike proteins, confocal microscopy was used to explore whether cleavage efficiency was related to differences in sub cellular localisation. We analysed the distribution of Spike in cells infected with Omicron and Delta SARS-COV-2 live virus isolates (Figure 2h-j); no clear differences were observed between the variants in distance from the nucleus or co-localisation with a golgi marker, indicating the subcellular localisation between Delta and Omicron is similar.

### Omicron spike is compromised in ability to mediate entry in cells expressing TMPRSS2

SARS-CoV-2 spike is a major determinant of viral infectivity and mediates cell entry via interaction with ACE2^8,17^ and either TMPRSS2^17^ at the plasma membrane or cathepsins in endosomes. The plasma membrane route of entry, and indeed transmissibility in animal models, is critically dependent on the polybasic cleavage site (PBCS) between S1 and S2^4,18-20^ and cleavage of spike prior to virion release from producer cells; this contrasts with the endosomal entry route, which does not require spike cleavage in producer cells ^4,5,21^. Plasma membrane fusion additionally allows the virus to avoid restriction factors in endosomes^4^.

We tested viral entry of WT Wuhan-1 D614G, Delta and Omicron spikes (Figure 3a) using the PV system. We first probed PV virions for spike protein and on western blotting noted reduced Omicron spike incorporation into virions compared to Delta (Supplementary Figure 6). We also noted that the Omicron spike was less cleaved than Delta as observed with live virus (Supplementary Figure 6). Of note cleavage of Omicron spike in cells was also lower compared to Delta and WT, again reminiscent of that in live virus (Figure 2f,j).

**Figure 3:**
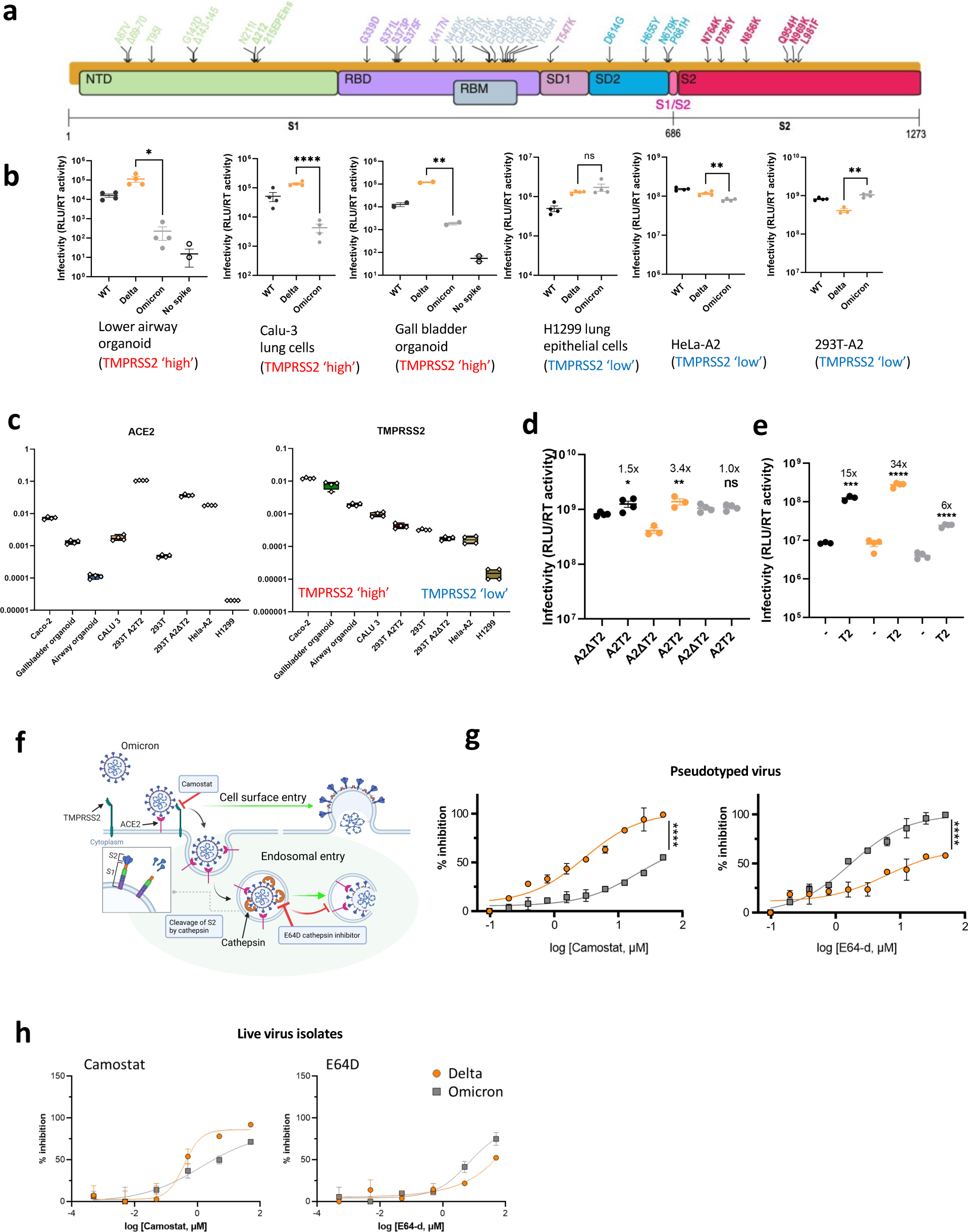
SARS-CoV-2 Omicron Variant spike enters cells less efficiently by TMPRSS2 mediated plasma membrane fusion. **a**. Graphical representation of Omicron spike mutations present in expression plasmid used to generate lentiviral pseudotyped virus (PV). Mutations coloured according to location in spike; bold mutations are novel to this lineage and have not been identified in previous variants of concern (VOCs). **b**. PV entry in airway organoids, Calu-3 lung cells, gall bladded organoids, H1299 lung cells, HeLa-ACE2 overexpressing cells and HEK 293-ACE2 overexpressing cells. Black – WT Wuhan-1 D614G, Blue – Delta B.1.617.2, Green Omicron BA.1, **c**. mRNA transcripts for ACE2 and TMPRSS2 in indicated cell types and organoids as measured by qPCR. Samples were run in quadruplicate. **d**. Entry of PV expressing spike in 293T cells transduced to overexpress ACE2 and (i) depleted for TMPRSS2 (A2Δ T2) or (ii) overexpressing TMPRSS2 (A2T2). **e**. Entry of PV expressing spike in 293T cells with endogenous (-) or overexpressed TMPRSS2 (T2). **f**. illustration of two cell entry pathways known to be used by SARS-CoV-2. **g**. Titration of inhibitors in A549-ACE2-TMPRSS2 (A549A2T2) cells using PV expressing Delta (orange) or Omicron (grey) spike in the presence of the indicated doses of Camostat or E64d, then analysed after 48 hours. % inhibition was calculated relative to the maximum luminescent signal for each condition. For each variant and dilution, mean ± SEM is shown for an experiment conducted in duplicate. Data are representative of two independent experiments **h**. Titration of inhibitors in A549-ACE2-TMPRSS2 (A549A2T2)-based luminescent reporter cells using live virus. Cells were infected at MOI = 0.01 with Delta (orange) or Omicron (grey) variants in the presence of the indicated doses of Camostat or E64d, then analysed after 24 hours. % inhibition was calculated relative to the maximum luminescent signal for each condition. For each variant and dilution, mean ± SEM is shown for an experiment conducted in triplicate. *p<0.05, **p<0.01, ****p<0.0001. Data are representative of two independent experiments

We infected primary 3D lower airway organoids and gall bladder organoids ^22,23^ (Figure 3b, Supplementary Figure 6), in addition to Calu-3 lung cells. Mirroring the lower replication of Omicron relative to Delta in live virus assays, we observed impaired entry efficiency for Omicron spike in both organoid systems and the Calu-3 cells in comparison to Delta and Wuhan-1 D614G wild type. By contrast, in H1299 lung cancer epithelial cells, Hela-ACE2 (overexpressing ACE2), and 293T cells overexpressing ACE2 and deleted for TMPRSS2 (293T-A2ΔT2), we did not observe large differences in entry efficiency for Delta and Omicron (Figure 2b).

In order to further explore our PV entry and infection findings, we studied the expression of ACE2 and TMPRSS2 across our target cells by qPCR on RNA extracts from cell lysates (Figure 3c). We found greater TMPRSS2 mRNA in cells where Omicron PV entry was impaired relative to Delta: for example Calu-3 and Organoids were higher in TMPRSS2 compared to H1299, Hela-ACE2, and 293T-A2ΔT2. ACE2 levels were variable as expected, and did not appear to have a correlation with differences in infection between Omicron and Delta PV (Figure 3b,c). We therefore hypothesised that the Omicron PV was disadvantaged relative to Delta in TMPRSS2-expressing cells.

To experimentally demonstrate differential usage of TMPRSS2 as a cofactor for virus entry by Omicron, we directly compared viral titres between 293T-A2ΔT2 and 293T-A2T2 cells; enhanced infectivity was observed from both WT and Delta when TMPRSS2 was overexpressed, suggesting TMPRSS2 is additionally required for their optimal virus entry. In contrast, no difference was observed from Omicron PV indicating Omicron is inefficient in utilising TMPRSS2 for its entry (Figure 3d). We hypothesised that the modest degree of TMPRSS2 dependent enhancement (3.4x) for Delta and a complete lack of enhancement for Omicron over a single infection round might relate to the overexpression of ACE2 in this system. To explore this possibility we tested standard 293T cells expressing low endogenous levels of ACE2 and those transduced to overexpress TMPRSS2 (293T-T2). Indeed we observed a much greater increase in infection for Delta (34x) under T2 overexpression conditions and a smaller increase (6x) for Omicron in the presence of overexpressed TMPRSS2 (Figure 3e). Together these data support a shift away from TMPRSS2 cofactor usage by Omicron spike, with modulation of effect size by ACE2.

A change in use of TMPRSS2 for entry would be predicted to alter the entry pathway (Figure 3f). To test the hypothesis of a change of entry route preference by Omicron spike away, we used inhibitors of proteases specific to either the endocytic pathway (ED64 blockade of cathepsins), or the plasma membrane pathway (camostat blockade of TMPRSS2). We reasoned that if indeed Omicron has become more dependent on the endocytic cell entry route it should be more sensitive to cathepsin inhibition. We infected A549 cells overexpressing ACE2 and TMPRSS2 (A549-A2T2) cells in the presence of serial dilutions of E64D or camostat; indeed E64D had a greater effect on cell entry by Omicron PV versus Delta PV and conversely camostat had a greater effect on Delta PV entry (Figure 3g). We then performed this experiment with live virus isolates, using an indicator lung cell line (A549-A2T2), with a similar, albeit less dramatic, result as shown for PV (Figure 3h). These drug data in both live virus and PV systems further support the shift in tropism away from TMPRSS2 expressing cells.

Having established that TMPRSS2 modulates entry mediated by Delta but not Omicron BA.1 spike, we sought to understand the distribution of TMPRSS2 and ACE2 expression in human respiratory cells. We used single-nuclei RNA-seq data from 5 locations in the human lung^24^. The comparison revealed higher expression of TMPRSS2 in the alveolar AT1 and AT2 pneumocytes, and in general lower expression in the trachea (upper airway, Supplementary Figure 7). ACE2 expression was generally lower overall, but appeared higher in AT1 and AT2 cells as compared to other cell types, and also in club cells^25^.

### Omicron Spike protein is unable to mediate cell-cell fusion

The ability of viral membrane glycoproteins to induce cell-cell fusion and syncytium formation is well established, providing an additional route for SARS-CoV-2 dissemination that may facilitate evasion of neutralising antibodies ^*2,18,27,28*^. The role of syncytium formation in viral replication and pathogenesis of severe COVID-19 has been reported and may be a druggable process to treat COVID-19 pathology^27,29^. SARS-CoV-2 mediated cell-cell fusion requires the polybasic cleavage site and spike cleavage at S1/2, and the process is known to be accelerated by the presence of TMPRSS2^30^.

Mutations at P681 in the PBCS have been observed in multiple SARS-CoV-2 lineages, most notably in the B.1.1.7 Alpha variant^31^ and the Delta/Kappa variants from the B.1.617 lineage^2,32^. We previously showed that spikes of these variants, all bearing P681 mutations, had significantly higher fusogenic potential than a D614G Wuhan-1 spike ^5^. Omicron bears P681H, in addition to 679 and 655 mutations (**Figure 3a**).

Given the requirement of TMPRSS2 for optimal cell-cell fusion, we hypothesised that Omicron may be impaired in mediating this process. We used a split GFP system^33^ to monitor cell-cell fusion in real time (**Figure 4a**). As a control to show spike cleavage is needed for fusion in our assay system, we titrated CMK (a furin inhibitor) into donor cell media prior to transfection with spike expressing plasmids^5^, observing dose dependent inhibition of fusion (supplementary figure 8). As a further control to demonstrate the need for ACE2 engagement by expressed spike, we tested the ability of convalescent serum containing neutralising antibodies to block cell-cell fusion^2^. Indeed serum blocked syncytia formation in a dose dependent manner (supplementary figure 8).

**Figure 4:**
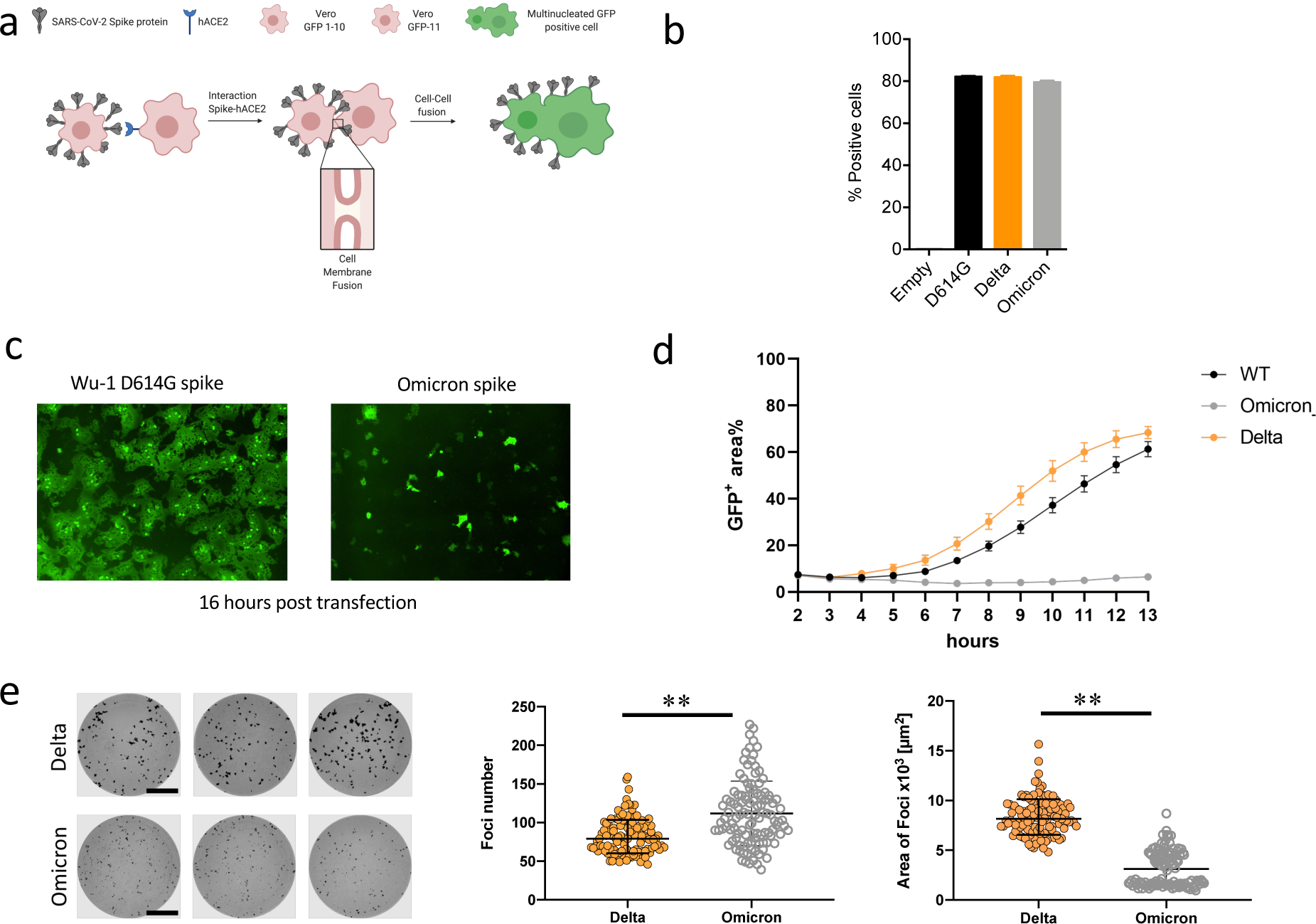
SARS-CoV-2 Omicron variant spike shows impaired cell-cell fusion activity and smaller infection foci generated by live virus. **a**. Schematic of cell-cell fusion assay. **b**. spike expression at the cell surface as determined by flow cytometry, showing % positive cells **c**. Reconstructed images at 16 hours of GFP+ syncytia. **d**. Quantification of cell-cell fusion kinetics showing percentage of green area to total cell area over time (WT is Wuhan-1 D614G). Mean is plotted with error bars representing SEM. Data are representative of at least two independent experiments. **e**. Left: representative images of H1299-ACE2 cell monolayers infected either with the live Delta (top 3 wells) or Omicron (bottom 3 wells) in a semi-solid media inhibitory for cell free infection. Cells were fixed 18-20 hours post-infection and stained for SARS-CoV-2 spike to visualize infection foci. Bar is 2mm. Middle and right panels: Quantified focus numbers and foci area (geometric mean and geometric std) for Omicron and Delta live virus infections. Data are from n=112 Delta infected wells and n=111 of Omicron infected wells from five independent experiments. ** p<0.0001 by the Wilcoxon sign-rank test.

We performed flow cytometry to check whether spike was expressed at the cell surface (Figure 4b). Indeed expression achieved similar levels as Delta and WT spikes. We proceeded to transfect spike bearing plasmids into 293T cells expressing GFP1-10 and Vero E6 cells stably expressing the GFP-11, so that the GFP signal could be detected upon cell-cell fusion and measured over time. (**Figure 4c,d**). We observed increased fusion for Delta as compared to D614G Wuhan-1 spike (**Figure 4d**). The Omicron spike however resulted in very poor fusion (**Figure 4d)**.

We predicted that poor fusion would impair cell-cell spread of Omicron. We therefore conducted an analysis of infection focus size using live virus infection of H1299 cells, in which we have showed similar entry efficiency for Omicron and Delta spike PV, see Figure 3b. Infection foci in spreading virus infections occurs because of localized (cell-to-cell) infection, likely facilitated by syncitium formation. The infection was performed in semi-solid media to inhibit cell-free infection, thereby favouring cell-cell infection. In the case of Omicron we predicted smaller foci due to impaired syncytium formation and less efficient direct cell-to-cell spread within a syncytium (independent of spike-ACE2 interactions). Indeed live Omicron infection resulted in slightly higher numbers of infection foci for a normalised input, but each infection focus was substantially smaller in size relative to those formed by the Delta variant (Figure 4e). An important caveat is that other viral and cellular factors could have contributed to the smaller foci in addition to an impairment of the cell-cell fusogenic potential of Omicron spike, for example ACE2 dependent cell-cell infection by budding virions across virological synapses, and ability of IFITMs to restrict fusion^30^ in the context of Omicron virus infection.

## Discussion

Here we have explored biological properties of Omicron from the perspectives of spike mediated immune evasion, ACE2 binding interactions, and cellular entry pathways that dictate tissue tropism. We have shown firstly that the Omicron spike confers very significant evasion of vaccine elicited neutralising antibodies that is more dramatic for ChAdOx-1 versus BNT162b2 vaccine sera. These data are supported by vaccine effectiveness measurements in the UK (UKHSA report Dec 2021). In longitudinally sampled participants, second dose waning was mitigated by third dose mRNA vaccination that also increased and broadened neutralisation of Omicron in the short term. This observation is critical for informing third dose vaccination efforts worldwide. Critically, ChAdOx-1 is widely used in low income settings where third doses with mRNA not widely available, and Omicron may contribute to higher infection rates and severe disease in such settings unless mRNA third doses can be implemented. Of further concern for low income countries was the absence of any neutralising activity to Omicron in sera after two doses of Coronavac in all of nine participants, in addition to poor Delta neutralisation.

In terms of implications for treatment options for moderate to severe clinical disease, we show high level loss of *in vitro* activity for the widely used mAb combination therapy REGN2 against Omicron, but no significant loss of activity of the polymerase inhibitors remdesivir and molnupiravir against live virus. The mAb sotrovimab has also been reported to retain significant *in vitro* activity against Omicron^34^. Importantly, the increased binding to ACE2 demonstrated here may itself increase the degree of nAb evasion under certain circumstances, by prolonging engagement duration with ACE2 and increasing probability of infection.

Despite the presence of three mutations that are individually predicted to favour spike S1/S2 cleavage, the observed cleavage spike in cells infected with live virus was lower compared to Delta and WT Wuhan-1 D614G. Similarly, live Omicron virions had a lower proportion of cleaved spike than Delta. These patterns of cleavage was also evident in PV particles. We previously noted that the proportion of cleaved spike was lower within cells as compared to virions^2^, suggesting that incorporation of spike into membranes that form virions may favour cleaved spike over full length spike.

Live Omicron virus showed a significant replication defect as compared to Delta cell culture systems where TMPRSS2 was present. Omicron spike was also associated with decreased entry relative to both WT and Delta spikes in target lower airway organoids, gallbladder organoids or Calu-3 lung cells expressing TMPRSS2, but no difference in cells where TMPRSS2 is low, for example H1299, Hela-ACE2 and 293T-A2. We proceeded to directly compare entry in ACE2 overexpressing cells where TMPRSS2 was either knocked down or overexpressed and showed that Omicron entry was not impacted by variations in TMPRSS2 expression, in stark contrast to WT and Delta spike PV entry. When ACE2 was not overexpressed, we importantly observed a large increase for Delta PV entry with TMPRSS2 availability, and a much smaller increase in entry for Omicron. These data suggest that TMPRSS2 cofactor usage is impacted by ACE2 levels, and we speculate that increased affinity for ACE2 is involved in Omicron’s shift away from TMPRSS2 usage. The hypothesis that Omicron spike has altered its preference to the cathepsin dependent endosomal route of entry (TMPRSS2-independent) was borne out by inhibitor experiments against cathepsin and TMPRSS2 in both PV and live virus isolates.

TMPRSS2 is a member of the TTSP family (TTSP – Type II Transmembran Serine Protease). This family comprises 17 members with diverse physiological functions. These proteins require activation and remain associated with membranes, and TMPRSS2 in particular can undergo autocatalytic activation^35^, though its role in human physiology is unclear. TMPRSS2 has been shown to be a cofactor for SARS-CoV-1 and MERS as well as SARS-CoV-2^17^.

Omicron presents an interesting parallel with SARS-CoV-1, which is not cleaved in the producer cell due to lack of the PBCS, but rather on the target cell surface by TMPRSS2 or by cathepsins in endosomes. SARS-CoV-1 is able to infect cells (i) via TMPRSS2 (and therefore syncytia can be induced^36^), or (ii) by endocytosis^20^, with the latter being favoured. SARS-CoV-2 appears to also favour endocytosis through reduced cleavage efficiency of spike impairing TMPRSS2 mediated entry. Indeed, the accumulated mutations in omicron include multiple additional basic amino acids leading to an increase in the overall charge of the S protein. This change may make S more sensitive to low-pH induced conformational changes, and be an adaptation that facilitates use of the low-pH endosomal entry route, and/or to entry in the lower pH environment in the upper airway.

Analysis of scRNA-seq datasets and qPCR measurements on our own human respiratory tissue samples suggest that lung cells have higher TMPRSS2 mRNA as compared to cells found in the upper airway. Recent findings from Hong Kong suggest lower replication in ex vivo lung tissue as compared to upper airway tissue for Omicron, but not for Delta (HKU website). A mechanism for Omicron’s poorer replication in the ex vivo tissue can be inferred from our findings regarding poor utilisation of TMPRSS2 dependent plasma membrane fusion by Omicron spike. Indeed recent data indicate lower virus burden in deep lung tissue and reduced pathogenicity in Omicron versus Delta infections using Syrian hamster models^37^.

As expected from suboptimal cleavage of spike in producer cells, and the fact that cell-cell fusion is more efficient in the presence of TMPRSS2, low fusogenic potential of the Omicron spike was observed when expressed in cells. This phenomenon could translate to impaired cell-cell spread, and indeed we observed smaller plaque sizes for Omicron compared to Delta in a system where cell free infection was limited by semi-solid medium.

Our observations highlight that Omicron has gained immune evasion properties whilst compromising cell entry in TMPRSS2 expressing cells such as those in alveoli, as well as compromising syncytia formation: a combination of traits consistent with reduced pathogenicity *in vivo*. Finally, experience with Omicron has demonstrated that predictions regarding replication and tropism based on sequence alone can be misleading, and detailed molecular understanding of the tropism shift has is vital as variants will inevitably continue to emerge.

## Limitations

Our neutralisation and infectivity assays tested only single isolates of each variant and ideally multiple isolates would be tested side by side. Our data showing tropism differences for Omicron in organoid systems and human nasal epithelial cultures are limited by the fact that they are in vitro systems, albeit using primary human tissue. In vivo studies are needed to understand the impact within an infected animal, ideally in the context of TMPRSS2 knock out. The fusion assay is limited by the use of transfected spike and not authentic virus. Although simulations predict a highly flexible and alternative linker 2 region conformation, these results are based on relatively short-timescale simulation studies and need further exploration.

## Methods

### Serum samples and ethical approval

Ethical approval for study of vaccine elicited antibodies in sera from vaccinees was obtained from the East of England – Cambridge Central Research Ethics Committee Cambridge (REC ref: 17/EE/0025). Studies involving health care workers (including testing and sequencing of respiratory samples) were reviewed and approved by The Institutional Human Ethics Committees of NCDC and CSIR-IGIB(NCDC/2020/NERC/14 and CSIR-IGIB/IHEC/2020-21/01). Participants provided informed consent.

The virus isolation procedures in this study were approved by the Institutional Review Board of National Institute for Infectious Diseases (approval ID: 1178) and Tokyo Metropolitan Institute of Public Health (approval ID: 3KenKenKen-466) according to the Declaration of Helsinki 2013. All protocols involving specimens from human subjects recruited at Kyoto University, Kuramochi Clinic Interpark and Universidad San Francisco de Quito were reviewed and approved by the Institutional Review Boards of Kyoto University (approval ID: G0697), Kuramochi Clinic Interpark (approval ID: G2021-004) and Universidad San Francisco de Quito (approval ID: CEISH P2020-022IN), and the Ecuadorian Ministry of Health (approval IDs: MSP-CGDES-2020-0121-O and MSP-CGDES-061-2020). The export of sera from Ecuador to Japan was approved by ARCSA ID: ARCSA-ARCSA-CGTC-DTRSNSOYA-2021-1626-M. All human subjects provided written informed consent. All protocols for the use of human specimens were reviewed and approved by the Institutional Review Boards of The Institute of Medical Science, The University of Tokyo (approval ID: 2021-1-0416), Kumamoto University (approval IDs: 2066 and 2074), University of Miyazaki (approval ID: O-1021)

### Sequencing

Spike genomes for the original Wuhan strain, and Omicron VOC were obtained from GISAID EpiCoV database accessed on 30^th^ November 2021. A consensus Spike genome was created from all complete and high coverage genomes, excluding all sequences with >5% Ns using Geneious Prime v2022. The consensus genome was translated to poly-proteins and the Spike gene was aligned to the original Wuhan strain using mafft v7.490^38^ with the --globalpair --maxiterate 1000 flags.

## Data availability

The above structural models used in this study for Delta and Omicron variants are available at https://github.com/CSB-Thukral-Lab/Spike_structural_models_Delta_and_Omicron

3D structural models of the spike homotrimer protein complex were alternatively also generated using Alphafold v2.1.1^39^. In its validation at the 14th edition of the Critical Assessment of protein Structure Prediction (CASP14) the predictions generated were demonstrated to be comparative to experimental structures. When a close homolog structure is available to Alphafold the predictions it generates for those positions are within typical experimental error. Required databases were downloaded on 02/12/2021. The program was run with the parameters --max_template_date=2021-12-01 --model_preset=monomer --db_preset=full_dbs --is_prokaryote_list=false. Protein structures were visualised in ChimeraX v1.3^40^. As predicted structures for the whole spike protein include poorly resolved chains at the terminal ends, these residues were identified by overlaying structures on PDB entry 6ZP2, then selected and removed from PDB files using the delete atoms/bonds action. Two further monomers were overlayed on 6zp2 to generate a homotrimer structure. Mutated residues were then coloured in red and labelled manually with respect to the Wuhan strain.

To model interactions between the omicron spike RBD and REGN 10933 and 10987, 6XDG was downloaded from PDB and aligned to the alphafolded omicron spike using the matchermaker function in ChimeraX v1.3. Predicted hydrogen bonds were plot within ChimeraX using a relaxed distance of 0.400Å and are shown as dashed green and blue lines. Predicted contact points between the RBD and REGN 10933 and 10987chains are indicated as red spheres using the same criteria as for hydrogen bonds.

## Data availability

All protein structures shown are freely available at github.com/ojcharles/viral_alphafold

ACE2 binding affinity assays

## Biolayer Interferometry

Assays were performed on an Octet Red (ForteBio) instrument at 30°C with shaking at 1,000 RPM. Streptavidin (SA) biosensors were hydrated in water for 10 min prior to a 1 min incubation in undiluted 10X kinetics buffer. SARS-CoV-2 Wuhan, Alpha, or Omicron RBDs were loaded at 2.5-7.5 μg/mL in 10X Kinetics Buffer for 100-400 s prior to baseline equilibration for 120 s in 10X kinetics buffer. Association of ACE2 to Wuhan RBD was assessed in 10X kinetics buffer at various concentrations in a three-fold dilution series from 333.3 to 4.1 nM was carried out for 150 s prior to dissociation for 150 s for. Association of ACE2 to Alpha or Omicron RBD was assessed in 10X kinetics buffer at various concentrations in a three-fold dilution series from 266.7 nM to 3.3 nM was carried out for 200 s prior to dissociation for 200 s. The data were normalized to the baseline and nonspecific binding was subtracted, prior to fitting performed using a 1:1 binding model and the ForteBio data analysis software. Kinetics values (KD, k_on_, k_off_) were determined with a global fit applied to all data.

The SARS-CoV-2-RBD-Avi constructs were synthesized by GenScript into pcDNA3.1-with an N-terminal mu-phosphatase signal peptide and a C-terminal octa-histidine tag, flexible linker, and avi tag (GHHHHHHHHGGSSGLNDIFEAQKIEWHE). The boundaries of the construct are N-328RFPN331 and 528KKST531-C, PMID: 32155444, PMID: 33706364. The human angiotensin-converting enzyme ectodomain (hACE2) consists of residues 1–614 fused to a C-terminal octa-histidine tag PMID: 32841599. Proteins were produced in Expi293F Cells (ThermoFisher Scientific) grown in suspension using Expi293 Expression Medium (ThermoFisher Scientific) at 37°C in a humidified 8% CO2 incubator rotating at 130 rpm. Cells grown to a density of 3 million cells per mL were transfected using the ExpiFectamine 293 Transfection Kit (ThermoFisher Scientific) and cultivated for 4 days. Proteins were purified from clarified supernatants using a nickel HisTrap HP affinity column (Cytiva) and washed with ten column volumes of 20 mM imidazole, 25 mM sodium phosphate pH 8.0, and 300 mM NaCl before elution on a gradient to 500 mM imidazole. Proteins were buffer exchanged into 20 mM sodium phosphate pH 8 and 100 mM NaCl and concentrated using centrifugal filters (Amicon Ultra) before being flash frozen.

### Flow Cytometry analysis of Spike proteins binding to recombinant human ACE2

ExpiCHO cells were seeded into 50 ml Mini Bioreactor Centrifuge Tube (Corning) at 6 × 10^6^ cells/ml in 5 ml ExpiCHO Expression medium (Life technology). Plasmids encoding SARS-CoV-2 Wuhan, Alpha (B.1.1.7), Delta (B.1.617.2) or Omicron (B.1.1.529) Spike protein (5 mg) were diluted in OptiPRO SFM (Life Technology) and mixed with Expifectamine CHO Reagent (Life Technology). After 1 min incubation at room temperature, transfection mixes were added to cell suspensions. Next, cells were incubated at 37°C 8% CO_2_ with an orbital shaking speed of 120 rpm for the following 48 hours.

Transiently transfected cells were harvested and washed with PBS, 1% BSA, 2 mM EDTA. Cells were counted, dispensed into round-bottom 96 well plates (Corning) and incubated with human IgG Fc-conjugated ACE2 serial dilutions (concentration range: 30’000 – 0.17 ng/ml) for 45 min at 4°C. After two washes, Alexa Fluor647 Goat Anti-Human IgG secondary Ab (1.5 mg/ml) (Jackson Immunoresearch) was added to the cells, which were then incubated for 30 min at 4°C in the dark. Cells were washed twice and resuspended in wash buffer for data acquisition at ZE5 cytometer (Biorad). To assess Spike protein expression level, an aliquot of each transfectant cell line was stained with 10 mg/ml of S2P6 antibody (Pinto et al., Science 2021) for 45 min at 4°C.

### Neutralisation titre analyses

Neutralisation by vaccine-elicited antibodies after two doses of the BNT162b2 and Chad-Ox-1 vaccine, in addition to a third dose with BNT162b2 was determined by PV infections in the presence of serial dilutions of sera as described^10^. The ID50 within groups were summarised as a geometric mean titre (GMT) and statistical comparison between groups were made with Mann-Whitney or Wilcoxon ranked sign tests. Statistical analyses were performed using Stata v13 and Prism v9.

#### Pseudotype virus (PV) experiments for neutralisation assays

HEK 293T CRL-3216, Hela-ACE-2 (Gift from James Voss), Vero CCL-81 were maintained in Dulbecco’s Modified Eagle Medium (DMEM) supplemented with 10% fetal calf serum (FCS), 100 U/ml penicillin, and 100mg/ml streptomycin. All cells were regularly tested and are mycoplasma free.

### Pseudotype virus preparation for testing against vaccine elicited antibodies and cell entry

Plasmids encoding the spike protein of SARS-CoV-2 D614 with a C terminal 19 amino acid deletion with D614G were used. Omicron and Delta spikes were generated by gene synthesis. Viral vectors were prepared by transfection of 293T cells by using Fugene HD transfection reagent (Promega). 293T cells were transfected with a mixture of 11ul of Fugene HD, 1μg of pCDNAΔ19 spike, 1ug of p8.91 HIV-1 gag-pol expression vector and 1.5μg of pCSFLW (expressing the firefly luciferase reporter gene with the HIV-1 packaging signal). Viral supernatant was collected at 48 and 72h after transfection, filtered through 0.45um filter and stored at -80°C as previously described. Infectivity was measured by luciferase detection in target cells.

### Sera Collection for live virus neutralisation experiments

Vaccine sera were collected from fourteen vaccinees four weeks after the second vaccination with BNT162b2 (Pfizer-BioNTech) (average age: 46, range: 38-55, 21% male). The sera obtained from seven vaccinees 2-3 weeks after the second vaccination with ChAdOx1 (Oxford-AstraZeneca) (average age: 46, range: 35-54, 71% male) were purchased from BioIVT.

Convalescent sera were collected from ten vaccine-naive individuals who had been infected with Delta variant (AY.29) (average age: 46, range: 22-63, 70% male). To determine the SARS-CoV-2 variants infected, saliva were collected from COVID-19 patients during onset and RNA was extracted using a QIAamp viral RNA mini kit (Qiagen, Qiagen, Cat# 52906) according to the manufacturer’s protocol. The sample was subjected to whole genome sequencing based on a modified ARTIC Network protocol^41^, and the near full-length SARS-CoV-2 genome sequences were obtained. Alternatively, we also performed the capture-hybridization method. The RNA isolated from saliva was treated with DNase I (Takara, Cat# EN0521) and the sequencing library was prepared using Twist library preparation kit (Twist Bioscience, Cat# 101058). The capture-hybridization was conducted using xGen COVID-19 capture panel and xGen hybridization and wash kit (Integrated DNA Technologies, Cat# 1080578) according to the manufacturer’s instruction. Illumina sequencing was performed using MiSeq reagent kit v2 (300 cycles) and a MiSeq sequencer (Illumina). For the data analysis, trimmomatic-0.39 (reference^42^) was used to remove the adaptors and low-quality reads from the raw sequence data. The trimmed paired-end reads were aligned to the human genome hg38 using bowtie2 v2.3.4.3 (reference^43^) and unmapped reads were mapped to the original SARS-CoV-2 genome (strain Wuhan-Hu-1, GenBank accession no. NC_045512.2) using Bwa-mem2 (https://github.com/bwa-mem2/bwa-mem2). The PCR duplicates were removed by gencore v0.16.0 (reference ^44^) and a consensus sequence was obtained by IGV v2.10.2 (reference ^45^). The mutations detected and viral lineage were determined by using CoVsurver (https://corona.bii.a-star.edu.sg) and Pangolin COVID-19 lineage assigner (https://pangolin.cog-uk.io/). The twelve convalescent sera during early pandemic (until April 2020) (average age: 71, range: 52-92, 8% male) were purchased from RayBiotech. Sera were inactivated at 56°C for 30 min and stored at –80°C until use.

### Live virus experiments in Caco-2

Vero E6 (ATCC-CRL1586) and Caco-2 cells transduced with BVDV NPro (Caco-NPro) were cultured in Dulbecco’s modified Eagle’s medium (DMEM) supplemented with 10□% (v/v) FCS, non-essential amino acid (NEAA), glutamine and 100 U penicillin ml^−1^ and 100 μg streptomycin ml^−1^ at 37 °C with 5□% CO_2_. Propagated live infectious **SARS-CoV-2** viruses, lineages **B.1.1.617.2** (Delta) and **B.1.1.529** (Omicron) were kindly received from Professor Wendy Barclay (Imperial College London) and Dr Jonathan Brown (Imperial College London) as part of the work conducted by G2P-UK National Virology Consortium. The virus isolate matching Omicron variant, kindly donated by Gavin Screaton at Oxford University, was propagated on Vero-ACE2-TMPRSS2 (VAT) cells for 3 days until cytopathic effect was observed. Caco-2 live virus infections were conducted in containment level 3 facility at the Cambridge Institute Therapeutic Immunology and Infectious Disease (CITIID) under approved standard operating procedures and protocols.

Lentivirus-transduced Caco-2 Npro cells were prepared in 48-well plates and infected in biological triplicates with either SARS-CoV-2 Delta or Omicron variant at m.o.i. 1 or 0.1 TCID50 per cell in 250 μl DMEM containing 2% FBS for 2 h at 37 °C with 5% CO2. Unabsorbed inoculum was then removed, cells were washed and the media was replaced with DMEM containing 2% FBS. Virus infections were maintained up to 72 h. Cell culture supernatants were harvested at 0, 24, 48 and 72 h post infection (p.i.) where viral RNAs were quantitated by RT-qPCR and/or infectious virus units were titrated by TCID50.

RNA extractions. Viral RNA was extracted using guanidinium-based reagent containing 2% TX-100. SARS-CoV-2 gene specific primer probe against E gene was used to evaluate levels of viral RNA by RT-qPCR, with an addition of no-template control analysed routinely as negative control. The levels of SARS-CoV-2 mRNA were determined based on absolute quantitation against a standard curve generated using in vitro-transcribed RNA from fragment 11 clone kindly provided by Volker Thiel (University of Bern). Data were collected using a ViiA 7 Real-Time PCR System (Applied Biosystems).

TCID50 assay. Ten-fold serial dilutions of collected virus supernatants were prepared in DMEM culture media. Of these dilutions, 50 μl was inoculated onto monolayers of Vero E6 cells grown on 96-well plates and incubated at 37 °C in a 5□% CO2 incubator. Virus titres were collected at 72 hours p.i. and expressed as TCID50 ml−1 values by the Reed–Muench method (Reed & Muench, 1938).

### Standardisation of virus input by SYBR Green-based product-enhanced PCR assay (SG-PERT)

The reverse transcriptase activity of virus preparations was determined by qPCR using a SYBR Green-based product-enhanced PCR assay (SG-PERT) as previously described^46^. Briefly, 10-fold dilutions of virus supernatant were lysed in a 1:1 ratio in a 2x lysis solution (made up of 40% glycerol v/v 0.25% Triton X-100 v/v 100mM KCl, RNase inhibitor 0.8 U/ml, TrisHCL 100mM, buffered to pH7.4) for 10 minutes at room temperature.

Sample lysates (12 μl) were added to 13 μl of SYBR Green master mix (containing 0.5μM of MS2-RNA Fwd and Rev primers, 3.5pmol/ml of MS2-RNA, and 0.125U/μl of Ribolock RNAse inhibitor and cycled in a QuantStudio. Relative amounts of reverse transcriptase activity were determined as the rate of transcription of bacteriophage MS2 RNA, with absolute RT activity calculated by comparing the relative amounts of RT to an RT standard of known activity.

### Cells for replication experiments and entry assays

H1299 cells were a kind gift from Simon Cook. Calu-3 cells were a kind gift from Paul Lehner, A549 A2T2^47^ cells were a kind gift from Massimo Palmerini. Vero E6 Ace2/TMPRSS2 cells were a kind gift from Emma Thomson.

### Nasal epithelial tissue infection

Primary human nasal epithelial cells (Cat# EP02, Batch# MP0010) were purchased from Epithelix and maintained according to the manufacturer’s procedure. The infection experiment using primary human nasal epithelial cells was performed as previously described (Saito et al, Nature, 2021)^48^. Briefly, the working viruses were diluted with Opti-MEM (Thermo Fisher Scientific, Cat# 11058021). The diluted viruses (1,000 TCID50 in 100 μl) were inoculated onto the apical side of the culture and incubated at 37°C for 1 h. The inoculated viruses were removed and washed twice with Opti-MEM. To harvest the viruses on the apical side of the culture, 100 μl Opti-MEM was applied onto the apical side of the culture and incubated at 37°C for 10 min. The Opti-MEM applied was harvested and used for RT–qPCR to quantify the viral RNA copy number (see below).

### Western blot on infected cells and virions

VeroE6-ACE2/TMPRSS2 cells were infected with SARS-CoV-2 at MOI 1. At 24 and 48 h.p.i., cells and culture media were harvested. The culture media were centrifuged with supernatants collected. To pellet down SARS-CoV-2 in the supernatants, virus were centrifuged at 20,000□g for 2□h at 4□°C through a 20% sucrose cushion or by PEG6000 (Papa et al plos pathogen 2021), and virus pellets were resuspended in 1 x SDS sample buffer.

Western blotting on virions was performed as previously described^13^. For cell lysates, the harvested cells were washed and lysed in lysis buffer [25 mM HEPES (pH 7.2), 20% glycerol, 125 mM NaCl, 1% Nonidet P40 substitute (Nacalai Tesque, Cat# 18558-54), protease inhibitor cocktail (Nacalai Tesque, Cat# 03969-21)] and lysates were diluted with 2 × sample buffer [100 mM Tris-HCl (pH 6.8), 4% SDS, 12% β-mercaptoethanol, 20% glycerol, 0.05% bromophenol blue] and boiled for 10 m. Then, 10 μl samples were subjected to Western blotting. For protein detection, the following antibodies were used: mouse anti-SARS-CoV-2 S monoclonal antibody (clone 1A9, GeneTex, Cat# GTX632604, 1:10,000), rabbit anti-SARS-CoV-2 N monoclonal antibody (clone HL344, GeneTex, Cat# GTX635679, 1:5,000), mouse anti-alpha tubulin (TUBA) monoclonal antibody (clone DM1A, Sigma-Aldrich, Cat# T9026, 1:10,000), horseradish peroxidase (HRP)-conjugated donkey anti-rabbit IgG polyclonal antibody (Jackson ImmunoResearch, Cat# 711-035-152, 1:10,000) and HRP-conjugated donkey anti-mouse IgG polyclonal antibody (Jackson ImmunoResearch, Cat# 715-035-150, 1:10,000). Chemiluminescence was detected using ChemiDoc Touch Imaging System (Bio-Rad, Cat #170-6545).

### Analysis of single-nuclei and single-cell RNA sequencing datasets

Normalised and log-transformed expression values per cell were obtained from human lung single-nuclei and epithelial non-small cell lung carcinoma cell lines. scanpy 1.7.1 was used to process the data. For the tissue data^24^, only single-nuclei RNA-seq were selected. From the cancer cell lines study, only mock samples were selected. ACE2 and TMPRSS2 expression was plotted with violin plot function in scanpy.

### Plasmids for split GFP system to measure cell-cell fusion

pQCXIP□BSR□GFP11 and pQCXIP□GFP1□10 were from Yutaka Hata ^49^ Addgene plasmid #68716; http://n2t.net/addgene:68716; RRID:Addgene_68716 and Addgene plasmid #68715; http://n2t.net/addgene:68715; RRID:Addgene_68715)

### Generation of GFP1□10 or GFP11 lentiviral particles

Lentiviral particles were generated by co-transfection of 293T or Vero cells with pQCXIP□BSR□GFP11 or pQCXIP□GFP1□10 as previously described ^50^. Supernatant containing virus particles was harvested after 48 and 72 hours, 0.45 μm filtered, and used to infect 293T or Vero cells to generate stable cell lines. 293T and Vero cells were transduced to stably express GFP1□10 or GFP11 respectively and were selected with 2 μg/ml puromycin.

### Cell-cell fusion assay

Cell-cell fusion assays were carried out as previously described ^13,50,51^. Briefly, 293T GFP1-10 and Vero-GFP11 cells were seeded at 80% confluence in a 1:1 ration in 48 multiwell plate the day before. Cells. were co-transfected with 0.5 μg of spike expression plasmids in pCDNA3 using Fugene 6 and following the manufacturer’s instructions (Promega). Cell-cell fusion was measured using an Incucyte and determined as the proportion of green area to total phase area. Data were then analysed using Incucyte software analysis. Graphs were generated using Prism 8 software. To measure cell surface spike expression HEK293 cells were transfected with S expression plasmids and stained with rabbit anti-SARS-CoV-2 S S1/S2 polyclonal antibody (Thermo Fisher Scientific, Cat# PA5-112048, 1:100). Normal rabbit IgG (SouthernBiotech, Cat# 0111-01, 1:100) was used as a negative control, and APC-conjugated goat anti-rabbit IgG polyclonal antibody (Jackson ImmunoResearch, Cat# 111-136-144, 1:50) was used as a secondary antibody. Surface expression level of S proteins was analysed using FACS Canto II (BD Biosciences) and FlowJo software v10.7.1 (BD Biosciences). Gating strategy for flow cytometry is shown in Supplementary Figure 8.

### Cell Culture for live virus experiments

VeroE6/TMPRSS2 and Vero E6 ACE2/TMPRSS2 cells [an African green monkey (*Chlorocebus sabaeus*) kidney cell line; JCRB1819]^52^ were maintained in Dulbecco’s modified Eagle’s medium (low glucose) (Wako, Cat# 041-29775) containing 10% FBS, G418 (1 mg/ml; Nacalai Tesque, Cat# G8168-10ML) and 1% PS. Calu-3 cells (a human lung epithelial cell line; ATCC HTB-55) were maintained in Eagle’s minimum essential medium (Sigma-Aldrich, Cat# M4655-500ML) containing 10% FBS and 1% PS.

### SARS-CoV-2 live virus isolation, preparation and titration for hNEC experiments

To isolate an Omicron variant (BA.1 lineage, strain TY38-873; GISAID ID: EPI_ISL_7418017), saliva was collected from a traveller arrived at Japan, and quantitative RT-PCR testing for SARS-CoV-2 was performed in an airport quarantine station, Japan. The sample was subjected to whole genome sequencing based on a modified ARTIC Network protocol^41^, and the near full-length SARS-CoV-2 genome sequence (GISAID ID: EPI_ISL_6913953) was deposited in GISAID. Virus isolation was performed as previously described^52^. In brief, the saliva was inoculated into VeroE6/TMPRSS2 cells and cytopathic effect (CPE) was observed 4 days after inoculation. The supernatant was then harvested and stored at –80°C as an original virus (GISAID ID: EPI_ISL_7418017). After one more passage in VeroE6/TMPRSS2 cells, the virus was obtained from National Institute of Infectious Diseases, Japan. A D614G-bearing early pandemic isolate (B.1.1lineage, strain TKYE610670; GISAID ID: EPI_ISL_479681) and a Delta isolate (B.1.617.2 lineage, strain TKYTK1734; GISAID ID: EPI_ISL_2378732) were used in the previous study^3^.

Virus preparation and titration was performed as previously described^3,53^. To prepare the working virus stock, 100 μl of the seed virus was inoculated into VeroE6/TMPRSS2 cells (5 × 10^6^ cells in a T-75 flask). One hour after infection, the culture medium was replaced with Dulbecco’s modified Eagle’s medium (low glucose) (Wako, Cat# 041-29775) containing 2% FBS and 1% PS. At 3 days postinfection, the culture medium was harvested and centrifuged, and the supernatants were collected as the working virus stock.

The titre of the prepared working virus was measured as the 50% tissue culture infectious dose (TCID_50_). Briefly, one day prior to infection, VeroE6/TMPRSS2 cells (10,000 cells/well) were seeded into a 96-well plate. Serially diluted virus stocks were inoculated into the cells and incubated at 37°C for 4 days. The cells were observed under microscopy to judge the CPE appearance. The value of TCID_50_/ml was calculated with the Reed–Muench method as previously described^13^.

### SARS-CoV-2 Infection with live virus

One day prior to infection, cells (10,000 cells) were seeded into a 96-well plate. SARS-CoV-2 (1,000 TCID_50_ as measured with plaque assay on Vero cells) was inoculated and incubated at 37°C for 1 h. The infected cells were washed, and 180 μl of culture medium was added. The culture supernatant (10 μl) was harvested at the indicated time points and used for real-time RT-PCR to quantify viral RNA copy number (see below).

### Real-Time RT-PCR

Real-time RT-PCR was performed as previously described^3,53^. Briefly, 5 μl of culture supernatant was mixed with 5 μl of 2 × RNA lysis buffer [2% Triton X-100, 50 mM KCl, 100 mM Tris-HCl (pH 7.4), 40% glycerol, 0.8 U/μl recombinant RNase inhibitor (Takara, Cat# 2313B)] and incubated at room temperature for 10 min. RNase-free water (90 μl) was added, and the diluted sample (2.5 μl) was used as the template for real-time RT-PCR performed according to the manufacturer’s protocol using the One Step TB Green PrimeScript PLUS RT-PCR kit (Takara, Cat# RR096A) and the following primers: Forward *N*, 5’-AGC CTC TTC TCG TTC CTC ATC AC-3’; and Reverse *N*, 5’-CCG CCA TTG CCA GCC ATT C-3’. The viral RNA copy number was standardized with a SARS-CoV-2 direct detection RT-qPCR kit (Takara, Cat# RC300A). Fluorescent signals were acquired using a QuantStudio 3 Real-Time PCR system (Thermo Fisher Scientific), a CFX Connect Real-Time PCR Detection system (Bio-Rad), an Eco Real-Time PCR System (Illumina) or a 7500 Real Time PCR System (Applied Biosystems).

### Preparation of Monoclonal Antibodies

Casirivimab and Imdevimab were prepared as previously described^54^. To construct the plasmids expressing anti-SARS-CoV-2 monoclonal antibodies (Casirivimab and Imdevimab), the sequences of the variable regions of Casirivimab and Imdevimab were obtained from KEGG Drug Database (https://www.genome.jp/kegg/drug/) and were artificially synthesized (Fasmac). The obtained coding sequences of the variable regions of the heavy and light chains were cloned into the pCAGGS vector containing the sequences of the human immunoglobulin 1 and kappa constant region [kindly provided by Dr. Hisashi Arase (Osaka University, Japan)].

To prepare these monoclonal antibodies, the pCAGGS vectors containing the sequences encoding the immunoglobulin heavy and light chains were cotransfected into HEK293T cells using PEI Max (Polysciences, Cat# 24765-1). At 48 h posttransfection, the cell culture supernatants were harvested, and the antibodies were purified using NAb protein A plus spin kit (Thermo Fisher Scientific, Cat# 89948) according to the manufacturer’s protocol.

### Neutralisation Assay for monoclonal antibodies with live virus

One day prior to infection, VeroE6/TMPRSS2 (10,000 cells) were seeded into a 96-well plate. The monoclonal antibodies (Casirivimab, Imdevimab, or Casirivimab/Imdevimab) and the heat-inactivated human sera were serially diluted with DMEM supplemented with 10% FCS and 1% PS. The diluted antibodies and sera were incubated with SARS-CoV-2 (120 TCID_50_) at 37°C for 1 h. The viruses without antibodies or sera were included as controls. The mixture (containing the virus at 100 TCID_50_) was inoculated onto a monolayer of VeroE6/TMPRSS2 cells and incubated at 37°C for 1 h. Then, the cells were washed with DMEM and cultured in DMEM supplemented with 10% FCS and 1% PS. At 24 h postinfection, the culture supernatants were harvested and viral RNA was quantified by real-time RT-PCR (see above). The assay of each antibody or serum was performed in triplicate or quadruplicate, and the 50% neutralization titre was calculated using Prism 9 (GraphPad Software).

### Antiviral drug assay with live virus

One day prior to infection, HOS-ACE2-TMPRSS2 cells (10,000 cells) were seeded into a 96-well plate. The cells were infected with SARS-CoV-2 (100 TCID_50_) at 37°C for 1 h. Then, the cells were washed with DMEM and cultured in DMEM supplemented with 10% FCS and 1% PS and the serially diluted Remdesivir (Selleck, Cat# S8932) or beta-d-N4-hydroxycytidine (a derivative of Molnupiravir; Cell Signaling Technology, Cat# 81178S). At 24 h postinfection, the culture supernatants were harvested and viral RNA was quantified by real-time RT-PCR (see above). The assay of each compound was performed in quadruplicate, and the 50% neutralization titre was calculated using Prism 9 (GraphPad Software).

### Cytotoxicity Assay

The CPE of the Remdesivir and beta-d-N4-hydroxycytidine were tested using a cell counting kit-8 (Dojindo, Cat# CK04-11) according to the manufacturer’s instructions. One day prior to the assay, HOS-ACE2-TMPRSS2 cells (10,000 cells) were seeded into a 96-well plate. The cells were cultured with the serially diluted compound for 24 hours. The cell counting kit-8 solution (10 μl) was added to each well, and the cells were incubated at 37°C for 90 m. Absorbance was measured at 450 nm by GloMax explorer microplate reader (Promega).

### Entry Inhibitors in A549-ACE2-TMPRSS2-based luminescent reporter cells using live virus

Quantification of viral replication in A549-ACE2-TMPRSS2-based luminescent reporter cells was performed as previously described^55^ and Gerber et al., *manuscript in preparation*). In brief, A549-ACE2-TMPRSS2 (A54-9A2T2) reporter cells (clone E8) over-expressing Renilla luciferase (Rluc) and SARS-CoV-2 Papain-like protease-activatable circularly permuted firefly luciferase (FFluc) were seeded in flat-bottomed 96-well plates. The following morning, cells were treated with the indicated doses of Camostat or E64d for 1 h, then infected with Delta or Omicron variants of SARS-CoV-2 at MOI = 0.01. After 24 h, cells were lysed in Dual-Glo Luciferase Buffer (Promega) diluted 1:1 with PBS and 1% NP-40. Lysates were then transferred to opaque 96-well plates, and viral replication quantitated as the ratio of FFluc/Rluc luminescence measured using the Dual-Glo kit (Promega) according to the manufacturer’s instructions. % inhibition was calculated relative to the maximum luminescent signal for each condition, then analysed using the Sigmoidal, 4PL, X is log(concentration) function in GraphPad Prism.

### Immunofluorescence (IF) assays

HeLa-ACE2 cells were grown on 19 mm glass coverslips in 24-well plates and incubated for 24 h. Cells were fixed in 4% paraformaldehyde (Applichem GmbH, Germany), and permeabilised in PBS-T (0.1% v/v Triton X-100 in PBS) for 5 min. Cells were blocked in PBS-T containing 5% w/v BSA for 30 min and labelled with primary antibodies against SARS-CoV-2 Spike (GTX632604, 1:1000) or anti-GM130[EP892Y]-cis-Golgi (ab52649, 1:250) in PBS-T, 1% w/v BSA for 1 h at room temperature. After washing, cells were stained with fluorochrome-conjugated secondary antibodies in PBS-T (1:1000), 1% w/v BSA for 1 h at room temperature in the dark. Nuclei were counterstained with 4’,6’-diamidino-2-phenylindole dihydrochloride (DAPI) and phalloidin 647 (A22287,1:1000) (Invitrogen, Molecular Probes, Germany). Confocal images were acquired on a Leica TCS SP8 microscope. Post-acquisition analysis was performed using Leica LAS X or Fiji (v1.49) software. For distance analysis, the two-dimensional coordinates of the centroids of Spike proteins were calculated using the Analyze Particles module of Fiji (ImageJ). The distance of each particle to the edge of the nucleus, visualised using DAPI stain, was looked up using a Euclidean distance map computed with the Distance Transform module of Fiji and exported as a list of distance measurements via the Analyze Particle function. Colocalization of Spike with the Golgi were estimated through Pearson’s correlation coefficients (*R*) using the PSC colocalization plug-in (ImageJ, NIH) (2). *R* values between −1 (perfect negative correlation) and +1 (perfect positive correlation), with 0 corresponding to no correlation were assigned^56^. Colocalization analyses were performed on >20 cells.

### Analysis of size of infection foci

Vero E6 cells (ATCC CRL-1586, obtained from Cellonex in South Africa) were propagated in complete growth medium consisting of Dulbecco’s Modified Eagle Medium (DMEM) with 10% fetal bovine serum (Hyclone) containing 10mM of HEPES, 1mM sodium pyruvate, 2mM L-glutamine and 0.1mM nonessential amino acids (Sigma-Aldrich). Vero E6 cells were passaged every 3–4 days. H1299 cell lines were propagated in growth medium consisting of complete Roswell Park Memorial Institute (RPMI) 1640 medium with 10% fetal bovine serum containing 10mM of HEPES, 1mM sodium pyruvate, 2mM L-glutamine and 0.1mM nonessential amino acids. H1299 cells were passaged every second day. The H1299-E3 (H1299-ACE2, clone E3) cell line was derived from H1299 (CRL-5803) as described in our previous work^57,58^. To Isolate virus, ACE2-expressing H1299 cells were seeded at 4.5 × 105 cells in a 6 well plate well and incubated for 18–20 h. After one DPBS wash, the sub-confluent cell monolayer was inoculated with 500 μL universal transport medium diluted 1:1 with growth medium filtered through a 0.45-μm filter. Cells were incubated for 1 h. Wells were then filled with 3 mL complete growth medium. After 4 days of infection (completion of passage 1 (P1)), cells were trypsinized, centrifuged at 300 rcf for 3 min and resuspended in 4 mL growth medium. Then all infected cells were added to Vero E6 cells that had been seeded at 2 × 10^5^ cells per mL, 20mL total, 18–20 h earlier in a T75 flask for cell-to-cell infection. The coculture of ACE2-expressing H1299-E3 and Vero E6 cells was incubated for 1 h and the flask was then filled with 20 mL of complete growth medium and incubated for 4 days. The viral supernatant (passage 2 (P2) stock) was used for experiments.P2 stock was sequenced and confirmed Omicron with the following substitutions: E:T9I,M:D3G,M:Q19E,M:A63T,N:P13L,N:R203K,N:G204R,ORF1a:K856R,ORF1a:L2084I,ORF1a:A2710T,ORF1a:T3255I,ORF1a:P3395H,ORF1a:I3758V,ORF1b:P314L,ORF1b:I15 66V,ORF9b:P10S,S:A67V,S:T95I,S:Y145D,S:L212I,S:G339D,S:S371L,S:S373P,S:S375F,S :K417N,S:N440K,S:G446S,S:S477N,S:T478K,S:E484A,S:Q493R,S:G496S,S:Q498R,S:N50 1Y,S:Y505H,S:T547K,S:D614G,S:H655Y,S:N679K,S:P681H,S:N764K,S:D796Y,S:N856K, S:Q954H,S:N969K,S:L981F. Sequence was deposited in GISAID, accession: EPI_ISL_7886688.

## Supporting information

SUPPS

## Acknowledgments

RKG is supported by a Wellcome Trust Senior Fellowship in Clinical Science (WT108082AIA). IG is a Wellcome Senior Fellow (207498/Z/17/Z). This study was supported by the Cambridge NIHRB Biomedical Research Centre. I.A.T.M.F. is funded by a SANTHE award (DEL-15-006). We would like to thank Paul Lehner for Calu-3 cells. We would like to thank James Voss for HeLa ACE2 and Suzanne Rihn for the A549 cells. We thank Gavin Screaton for the Omicron isolate M21021166. We would like to thank Claire Cormie for microscope training. We would like to thank Thushan de Silva for the Delta variant isolate. This work was supported by the MRC (TSF ref. MR/T032413/1 to NJM), NHSBT (grant ref. WPA15-02 to NJM), Addenbrooke’s Charitable Trust (grant ref. to 900239 NJM). The authors acknowledge support from the G2P-UK National Virology consortium funded by MRC/UKRI (grant ref: MR/W005611/1). This study was also supported by The Rosetrees Trust and the Geno2pheno UK consortium. SF acknowledges the EPSRC (EP/V002910/1). KS is supported by AMED Research Program on Emerging and Re-emerging Infectious Diseases (20fk0108270 and 20fk0108413), JST SICORP (JPMJSC20U1 and JPMJSC21U5) and JST CREST (JPMJCR20H4). We would like to thank all members belonging to The Genotype to Phenotype Japan (G2P-Japan) Consortium. We thank National Institute for Infectious Diseases, Japan and Hisashi Arase (Osaka University, Japan) for providing virus isolates and reagents and Ituro Inoue and Sachiko Sakamoto (National Institute of Genetics, Japan) for technical supports. This study was supported in part by AMED Research Program on Emerging and Re-emerging Infectious Diseases 20fk0108163 (to Akatsuki Saito), 20fk0108146 (to Kei Sato), 20fk0108270 (to Kei Sato), 20fk0108413 (to Terumasa Ikeda and Kei Sato) and 20fk0108451 (to Akatsuki Saito, Terumasa Ikeda, Takamasa Ueno, Akifumi Takaori-Kondo, G2P-Japan Consortium and Kei Sato); AMED Research Program on HIV/AIDS 21fk0410033 (to Akatsuki Saito) and 21fk0410039 (to Kei Sato); AMED Japan Program for Infectious Diseases Research and Infrastructure 20wm0325009 (to Akatsuki Saito) and 21wm0325009 (to Akatsuki Saito); JST A-STEP JPMJTM20SL (to Terumasa Ikeda); JST SICORP (e-ASIA) JPMJSC20U1 (to Kei Sato); JST SICORP JPMJSC21U5 (to Kei Sato), JST CREST JPMJCR20H4 (to Kei Sato); JSPS KAKENHI Grant-in-Aid for Scientific Research C 19K06382 (to Akatsuki Saito), Scientific Research B 18H02662 (to Kei Sato) and 21H02737 (to Kei Sato); JSPS Fund for the Promotion of Joint International Research (Fostering Joint International Research) 18KK0447 (to Kei Sato); JSPS Core-to-Core Program JPJSCCA20190008 (A. Advanced Research Networks) (to Kei Sato); JSPS Research Fellow DC1 19J20488 (to Izumi Kimura); JSPS Leading Initiative for Excellent Young Researchers (LEADER) (to Terumasa Ikeda); Takeda Science Foundation (to Terumasa Ikeda); The Tokyo Biochemical Research Foundation (to Kei Sato); Mitsubishi Foundation (to Terumasa Ikeda); Shin-Nihon Foundation of Advanced Medical Research (to Terumasa Ikeda); a Grant for Joint Research Projects of the Research Institute for Microbial Diseases, Osaka University (to Akatsuki Saito); an intramural grant from Kumamoto University COVID-19 Research Projects (AMABIE) (to Terumasa Ikeda); Intercontinental Research and Educational Platform Aiming for Eradication of HIV/AIDS (to Terumasa Ikeda); and Joint Usage/Research Center program of Institute for Frontier Life and Medical Sciences, Kyoto University (to Kei Sato). This study was supported by the National Institute of Allergy and Infectious Diseases (DP1AI158186 and HHSN272201700059C to D.V.), a Pew Biomedical Scholars Award (D.V.), an Investigators in the Pathogenesis of Infectious Disease Awards from the Burroughs Wellcome Fund (D.V.), Fast Grants (D.V.) and the Bill and Melinda Gates Foundation (OPP1156262 to D.V.). D.V. is an Investigator of the Howard Hughes Medical Institute. T.B. is supported by an EASL Juan Rodès fellowship. F.S. was supported by a UKRI Future Leaders fellowship.

## The Genotype to Phenotype Japan (G2P-Japan) Consortium members

Ryoko Kawabata^16^, Nanami Morizako^16^, Kenji Sadamasu^17^, Hiroyuki Asakura^17^, Mami Nagashima^17^, Kazuhisa Yoshimura^17^, Jumpei Ito^18^, Izumi Kimura^18^, Keiya Uriu^18^, Yusuke Kosugi^18^, Mai Suganami^18^, Akiko Oide^18^, Miyabishara Yokoyama^18^, Mika Chiba^18^ , Akatsuki Saito^31^, Erika P Butlertanaka^31^, Yuri L Tanaka^31^, Terumasa Ikeda^32^, Chihiro Motozono^32^, Hesham Nasser^32^, Ryo Shimizu^32^, Yue Yuan^32^, Kazuko Kitazato^32^, Haruyo Hasebe^32^, So Nakagawa^33^, Jiaqi Wu^33^, Miyoko Takahashi^33^, Takasuke Fukuhara^34^, Kenta Shimizu^34^, Kana Tsushima^34^, Haruko Kubo^34^, Yasuhiro Kazuma^35^, Ryosuke Nomura^35^, Yoshihito Horisawa^35^, Kayoko Nagata^35^, Yugo Kawai^35^, Yohei Yanagida^35^, Yusuke Tashiro^35^, Kenzo Tokunaga^36^, Seiya Ozono^36^

^31^University of Miyazaki, Miyazaki, Japan

^32^Kumamoto University, Kumamoto, Japan

^33^Tokai University, Tokyo, Japan

^34^Hokkaido University, Sapporo, Japan

^35^Kyoto University, Kyoto, Japan

^36^National Institute of Infectious Diseases, Tokyo, Japan

## The CITIID-NIHR BioResource COVID-19 Collaboration

Stephen Baker^1,2^, Gordon Dougan^1,2^, Christoph Hess^2^, Nathalie Kingston^9^, Paul J. Lehner^1,2^, Paul A. Lyons^1,2^, Nicholas J. Matheson^1,2^, Willem H. Owehand^22^, Caroline Saunders^21^, Charlotte Summers^2^, James E.D. Thaventhiran^2^, Mark Toshner^2^, Michael P. Weekes^2^, Patrick Maxwell^37^, Ashley Shaw^37^ Ashlea Bucke^38^, Jo Calder^38^, Laura Canna^38^, Jason Domingo^38^, Anne Elmer^38^, Stewart Fuller^38^, Julie Harris^38^, Sarah Hewitt^38^, Jane Kennet^38^, Sherly Jose^38^, Jenny Kourampa^38^, Anne Meadows^38^, Criona O’Brien^38^, Jane Price^38^, Cherry Publico^38^, Rebecca Rastall^38^, Carla Ribeiro^38^, Jane Rowlands^38^, Valentina Ruffolo^38^, Hugo Tordesillas^38^, Ben Bullman^1^, Benjamin J. Dunmore^2^, Stuart Fawke^39^, Stefan Gräf ^2^, Josh Hodgson^3^, Christopher Huang^3^, Kelvin Hunter^2,^, Emma Jones^31^, Ekaterina Legchenko^2^, Cecilia Matara^2^, Jennifer Martin^2^, Federica Mescia^2^, Ciara O’Donnell^2^, Linda Pointon^2^, Nicole Pond^2^, Joy Shih^2^, Rachel Sutcliffe^2^, Tobias Tilly^2^, Carmen Treacy^2^, Zhen Tong^2^, Jennifer Wood^2^, Marta Wylot^2^, Laura Bergamaschi^2^, Ariana Betancourt^2^, Georgie Bower^2^, Chiara Cossetti^2,^, Aloka De Sa^2^, Madeline Epping^2^, Stuart Fawke^2^, Nick Gleadall^2^, Richard Grenfell^2^, Andrew Hinch^2^, Oisin Huhn^39^, Sarah Jackson^2^, Isobel Jarvis^2^, Ben Krishna^2^, Daniel Lewis^3^, Joe Marsden^3^, Francesca Nice^41^, Georgina Okecha^3^, Ommar Omarjee^2^, Marianne Perera^2^, Martin Potts^2^, Nathan Richoz^2^, Veronika Romashova^2^, Natalia Savinykh Yarkoni^3^, Rahul Sharma^3^, Luca Stefanucci^2^, Jonathan Stephens^22^, Mateusz Strezlecki^2^, Lori Turner^2^, Eckart M.D.D. De Bie^2^, Katherine Bunclark^2^, Masa Josipovic^2^, Michael Mackay^2^, Federica Mescia^2^, Alice Michael^27^, Sabrina Rossi^37^, Mayurun Selvan^3^, Sarah Spencer^15^, Cissy Yong^37^, John Allison^9^, Helen Butcher^9,40^, Daniela Caputo^9,40^, Debbie Clapham-Riley^9,40^, Eleanor Dewhurst^9,40^, Anita Furlong^9,40^, Barbara Graves^9,40^, Jennifer Gray^9,40^, Tasmin Ivers^9,40^, Mary Kasanicki^9,30^, Emma Le Gresley^9,40^, Rachel Linger^9,40^, Sarah Meloy^9,40^, Francesca Muldoon^9,40^, Nigel Ovington^9^, Sofia Papadia^9,40^, Isabel Phelan^9,40^, Hannah Stark^9,40^, Kathleen E Stirrups^22,12^, Paul Townsend^40^, Neil Walker^40^, Jennifer Webster^9,40^, Ingrid Scholtes^40^, Sabine Hein^40^, Rebecca King^40^

^37^ Cambridge University Hospitals NHS Trust, Cambridge UK.

^38^Cambridge Clinical Research Centre, NIHR Clinical Research Facility, Cambridge University Hospitals NHS Foundation Trust, Addenbrooke’s Hospital, Cambridge CB2 0QQ, UK

^39^Department of Biochemistry, University of Cambridge, Cambridge, CB2 1QW, UK

^40^University of Cambridge, Cambridge Biomedical Campus, Cambridge CB2 0QQ, UK

## Ecuador-COVID19 Consortium

Sully Márquez ^uuu^, Belén Prado-Vivar ^uuu^, Mónica Becerra-Wong ^uuu^, Mateo Caravajal ^uuu^,

Gabriel Trueba ^uuu^, Patricio Rojas-Silva ^uuu^,

Michelle Grunauer ^vvv^,

Bernardo Gutierrez ^www^, Juan José Guadalupe ^www^,

Juan Carlos Fernández-Cadena ^xxx^, Derly Andrade-Molina ^yyy^,

Manuel Baldeon ^zzz^,

Andrea Pinos ^ooo^

^uuu^ Universidad San Francisco de Quito, COCIBA, Instituto de Microbiología, Cumbaya EC1702, Ecuador

^vvv^ Universidad San Francisco de Quito, COCSA, Escuela de Medicina, Cumbaya EC1702, Ecuador

^www^ Universidad San Francisco de Quito, COCIBA, Laboratorio de Biotecnología Vegetal, Cumbaya EC1702, Ecuador

^xxx^ Laboratorio INTERLAB, Guayaquil EC09, Ecuador

^yyy^ Universidad Espíritu Santo, Laboratorio de Omicas, Guayaquil, EC09, Ecuador

^zzz^ Universidad Internacional del Ecuador, Facultad de Ciencias Médicas, de la Salud y la Vida, Quito EC1701, Ecuador

^ooo^ Centros Médicos Dr. Marco Albuja, Quito EC1701, Ecuador

## Notes

### Competing Interest Statement

RKG has received honoraria from GSK, Janssen and ViiV for educational activities

