## Supplementary material for "SARS-CoV-2 Omicron spike mediated immune escape and tropism shift": SUPPS

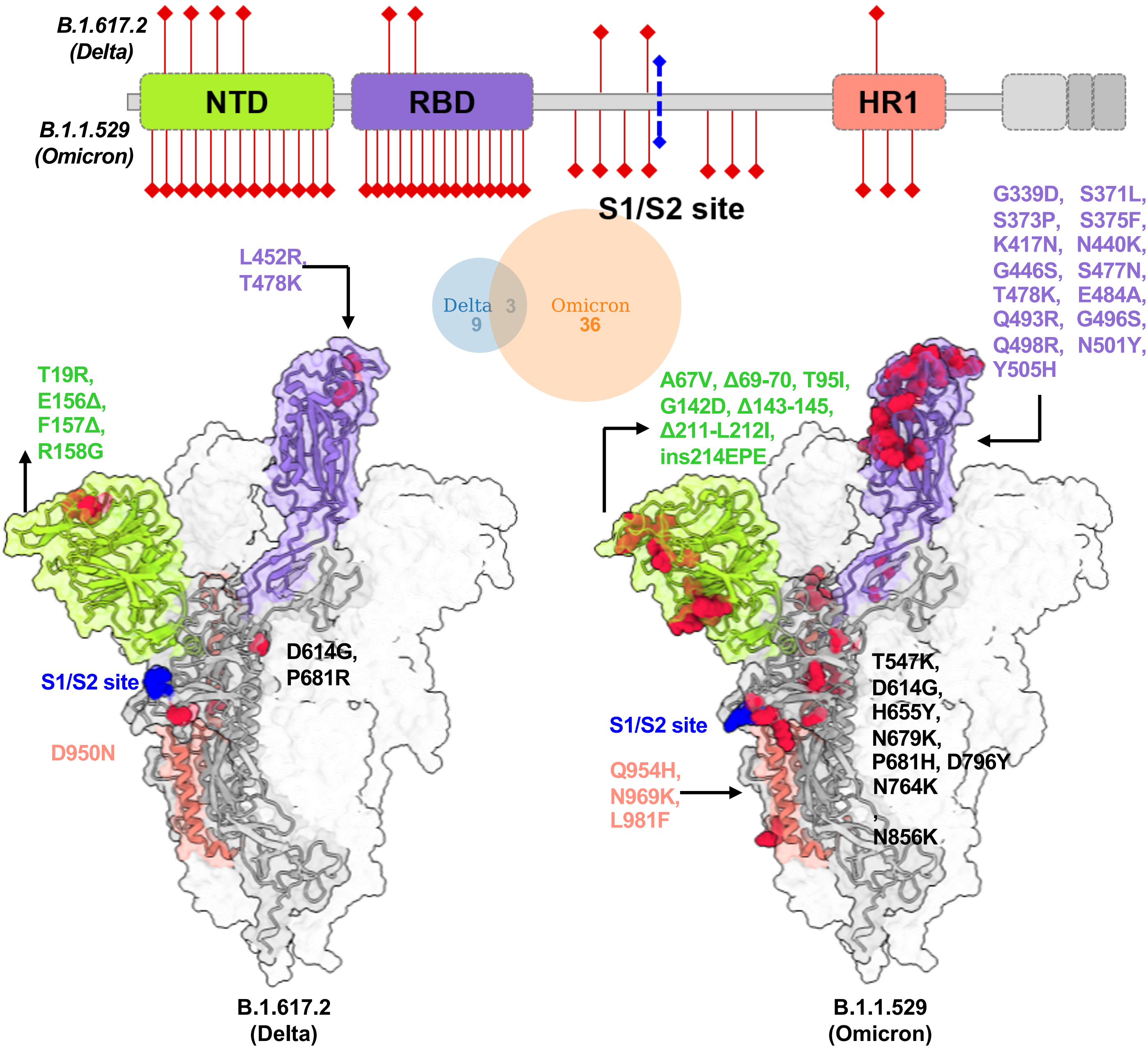


**Supplementary Figure 1: Structural model of SARS-CoV-2 Delta and Omicron spike variants and their corresponding protein intramolecular network.** Variant structures for Delta and Omicron of spike protein displaying mutational sites in red color. The schematic highlights differences in domain-wise mutations across protein length.

**a**

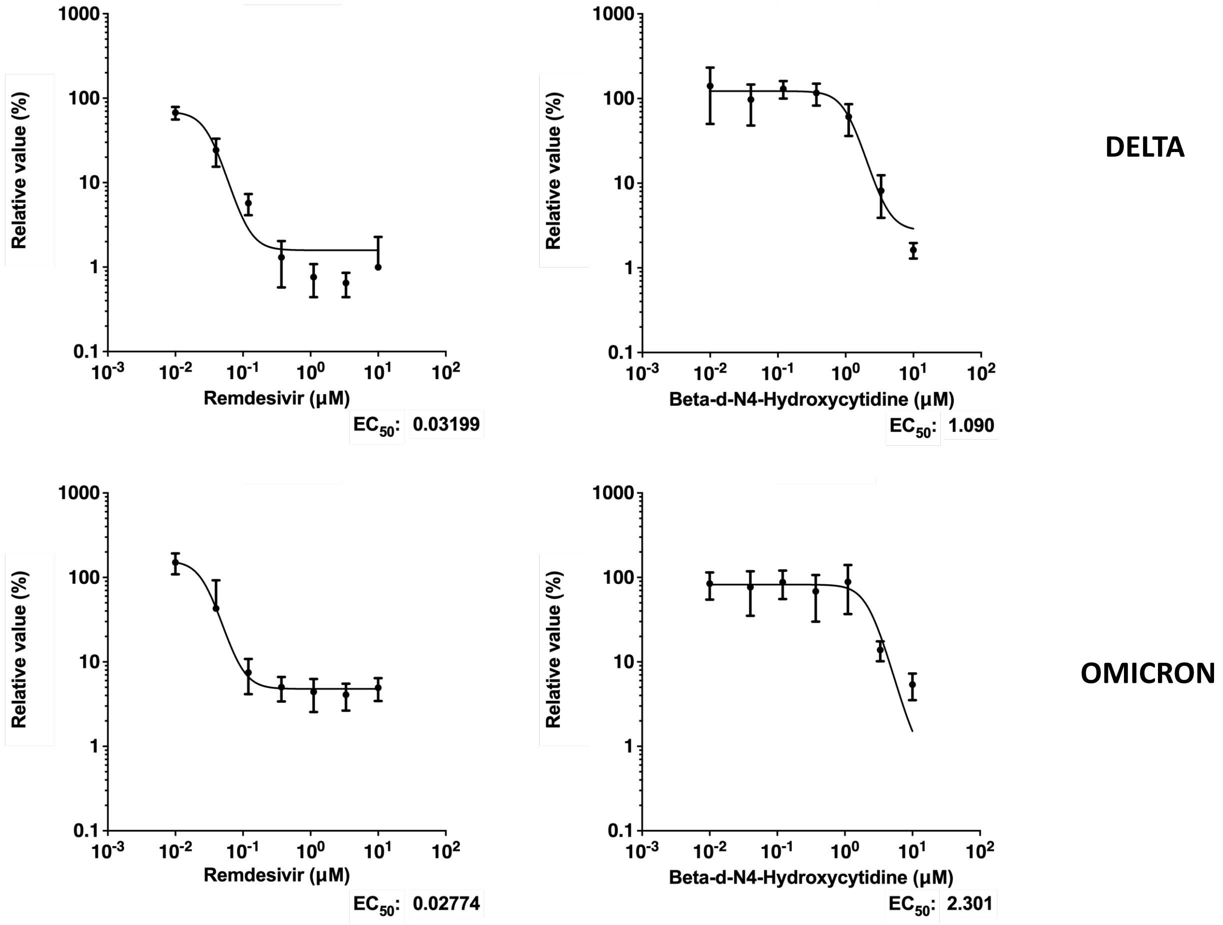


**b**

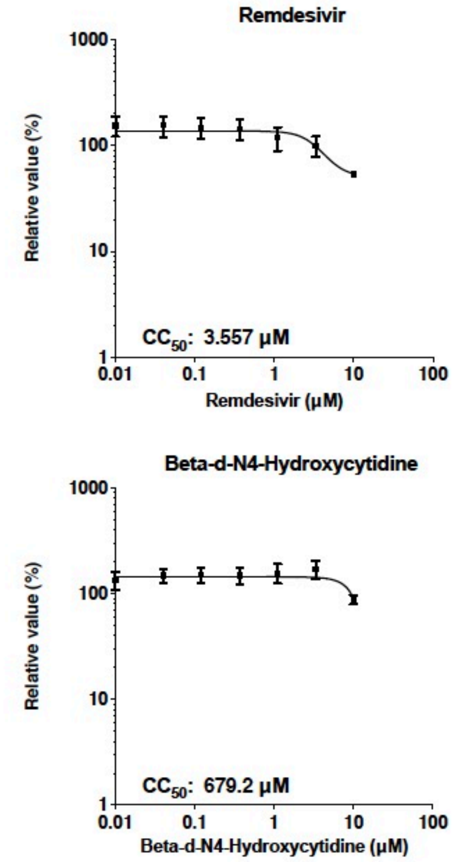


**Supplementary Figure 2** : **Sensitivity of replication competent SARS-CoV-2 Omicron and Delta variants to clinically approved direct acting antiviral molecules remdesivir and the active metabolite of molnupiravir.** HOS cells overexpressing ACE2 and TMPRSS2 were used and a viral input of 1000TCID50 was used (TCID50 measured using VeroE6/TMPRSS2 cells). **a.** Dose-response curves. Infection as measured by viral RNA copies relative to the no drug control (100%) is plotted on the y axis with serial drug dilution on the x axis. EC50 is indicated for each panel and calculated in GraphPad Prism. **b**. toxicity assay showing relative cell viability at a range of drug doses. Data are representative of two independent experiments.

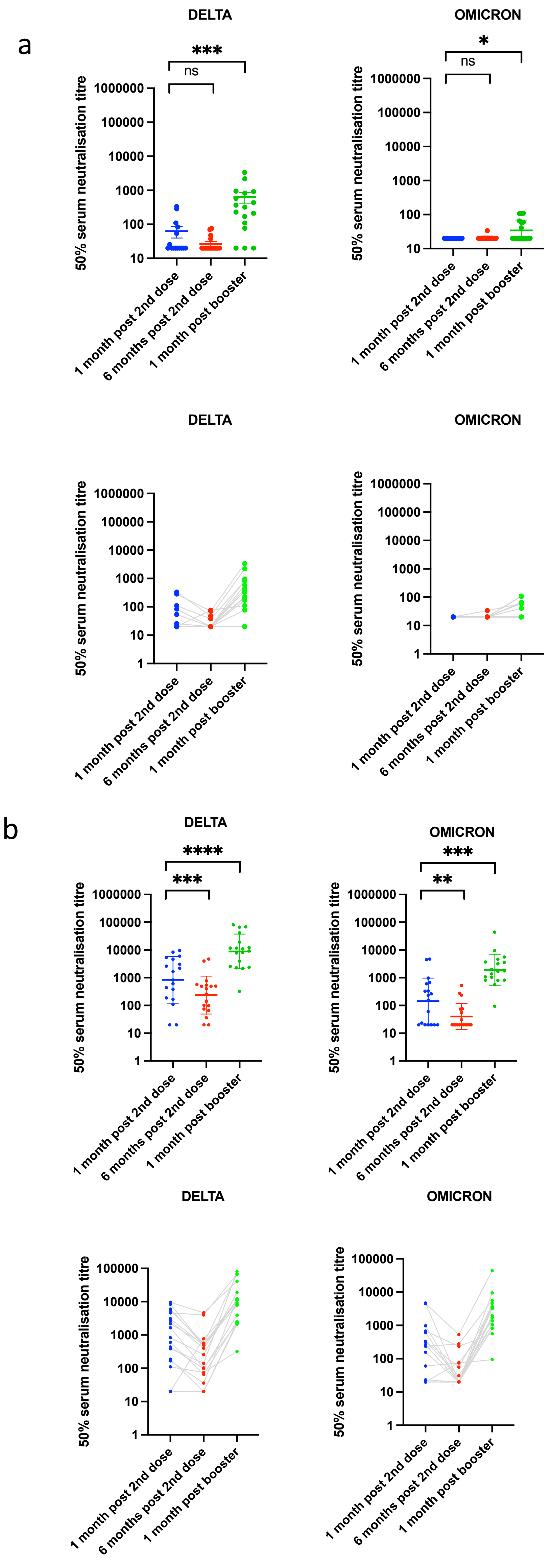


**Supplementary Figure 3: Neutralisation of spike pseudotyped virus by sera from vaccinated individuals over three time points following dose two (ChAdOx-1 or BNT162b2) and dose three (BNT162b2 only) a.** n=17 ChAdOx-1 or **b.** n=18 BNT12b2. GMT (geometric mean titre) with s.d are presented. Data representative of two independent experiments each with two technical replicates. **p<0.01, *** p<0.001, ****p<0.0001 Wilcoxon matched-pairs signed rank test, ns not significant. Data are for individuals who tested negative for anti-N IgG at each time point.

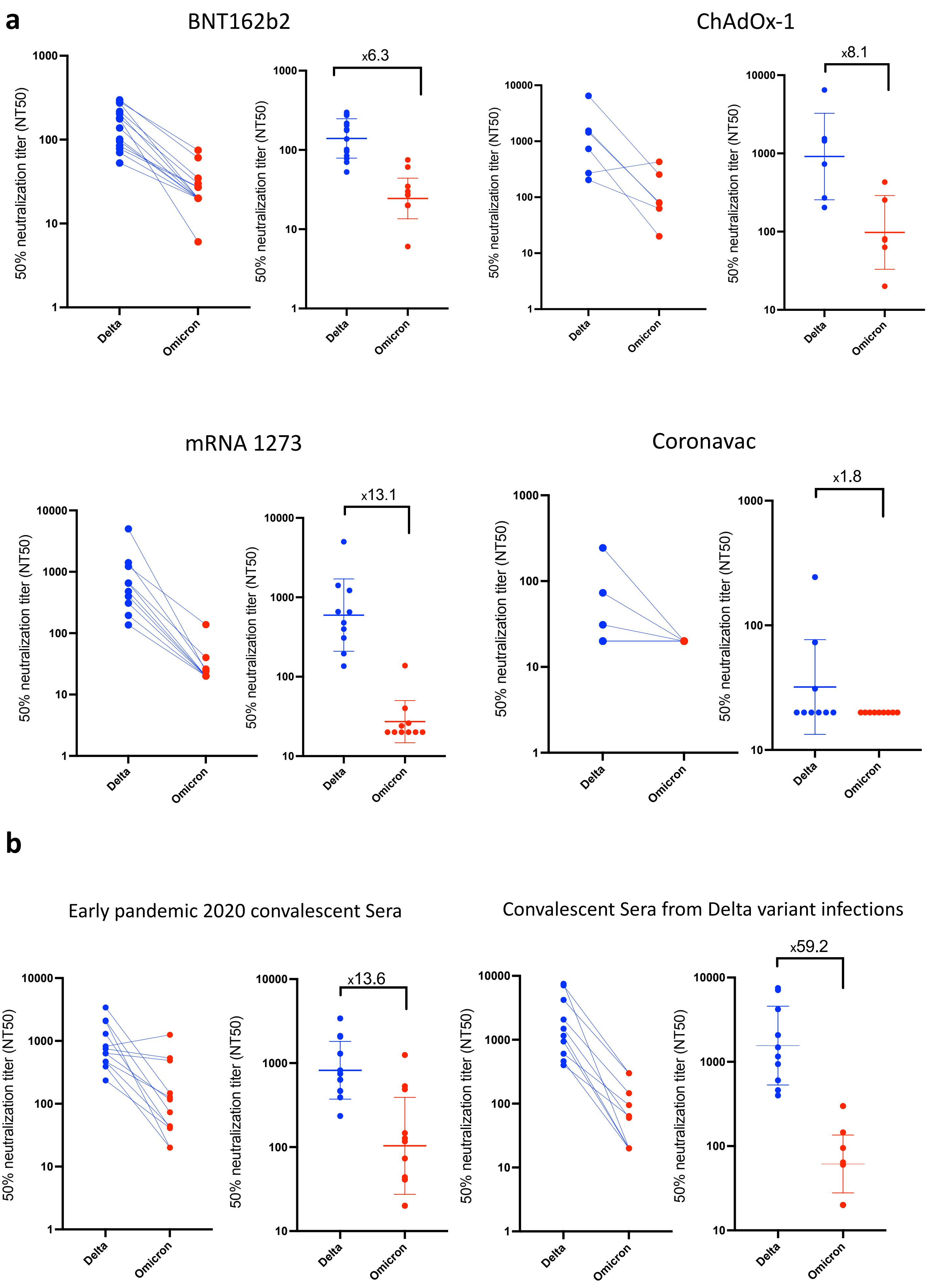


**Supplementary Figure 4: a.** Neutralisation of replication competent Delta and Omicron viruses by sera from vaccinated individuals following dose two of ChAdOx-1, BNT162b2, mRNA 1273, and Coronavac vaccines**. b.** neutralization of live viruses by sera derived from recovered individuals following infection in early 2020 or with confirmed Delta infection. Data are representative of two independent experiments. Fold change in NT50 is indicated.

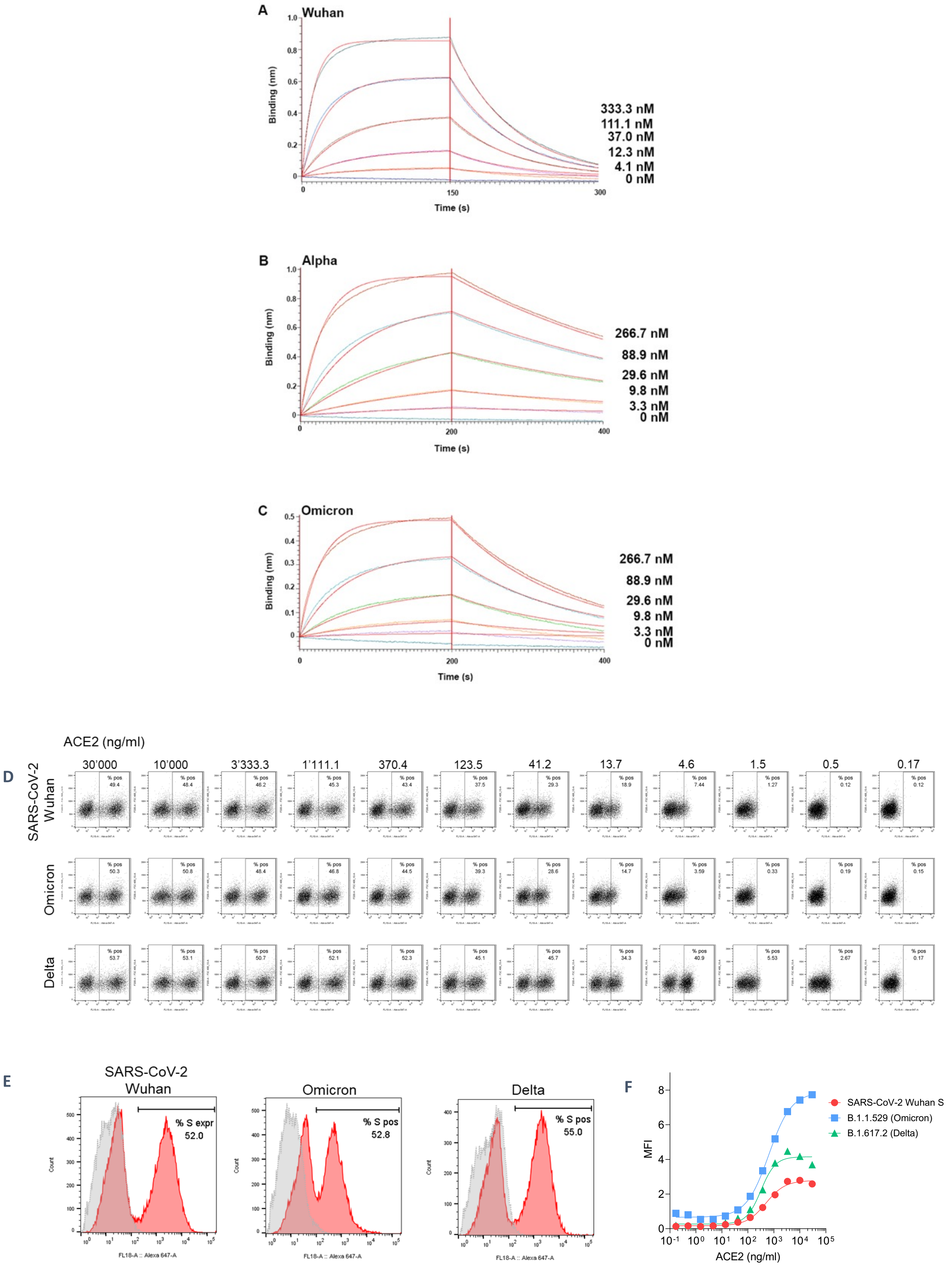


**Supplementary Figure 5**: **Omicron spike binding to ACE2:** (A-C) Biolayer interferometry binding analysis of human ACE2 to immobilized SARS-CoV-2 Wuhan RBD (A) Alpha RBD (B) or Omicron (C) VOC. Red lines correspond to a global fit of the data using a 1:1 binding model. (D-F). Flow cytometry analysis of ACE2 binding to SARS-CoV-2 Wuhan, B.1.617.2 (Delta) or Omicron (B.1.1.529) spike proteins transiently expressed in ExpiCHO cells. **E.** Expression level of each spike as measured via binding by S2-subunit targeting S2P6 mAb (Pinto et al., Science 2021). Grey histograms represent background florescence of cells stained with the secondary antibody only. **F** Normalized ACE2 binding results based on spike protein expression levels. Values on the Y axis indicate the ratio between the mean fluorescence intensity (MFI) of positive cells stained for ACE2 binding and the MFI of the S2P6 positive cells. Data are representative of one independent experiment out of two.

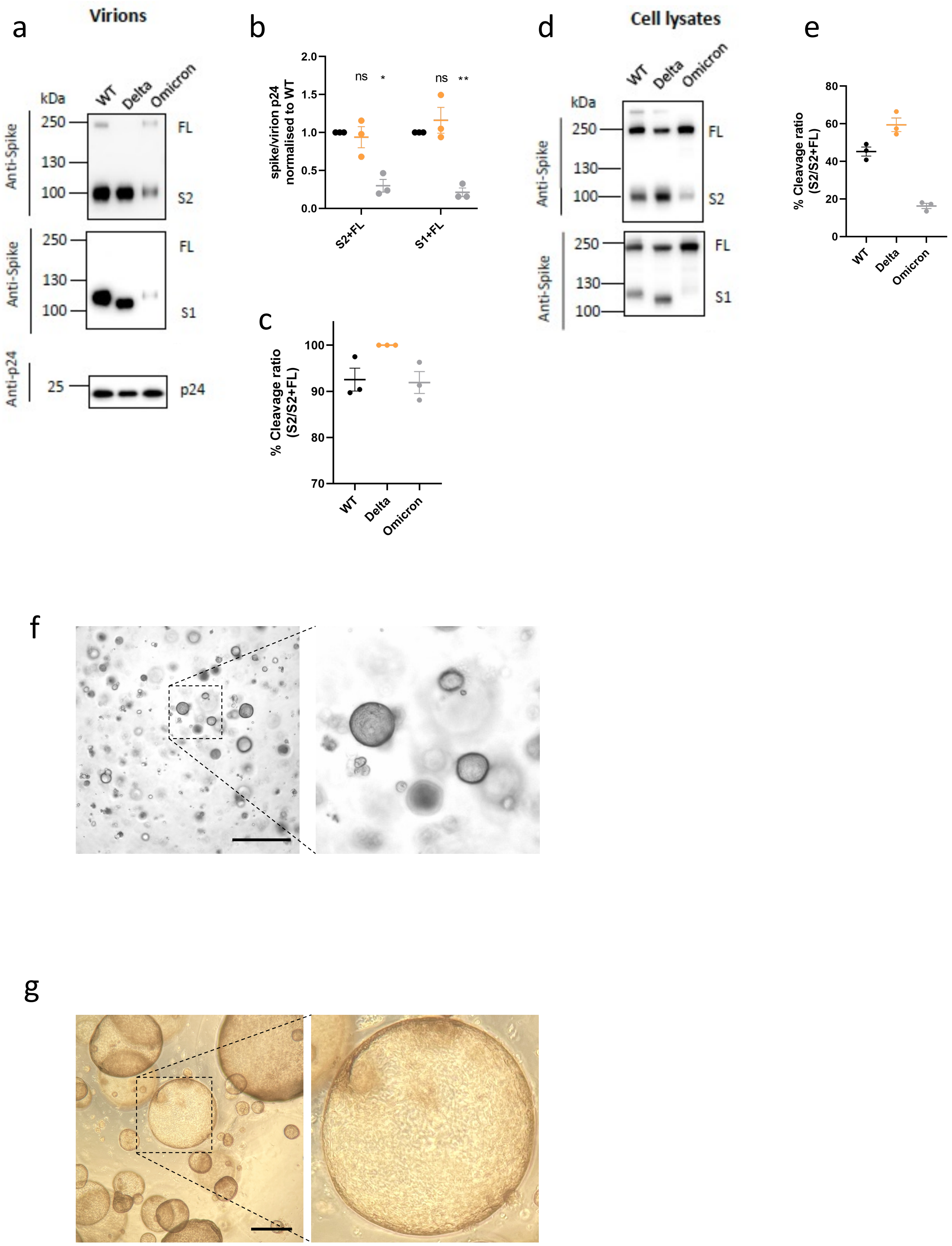


**Supplementary Figure 6: SARS-CoV-2 Omicron Variant spike pseudotyped lentiviruses a.** western blots of pseudotyped virus (PV) virions from 293T producer cells following transfection with plasmids expressing lentiviral vectors and SARS-CoV-2 S plasmids. (WT- Wuhan-1 with D614G), probed with antibodies for HIV-1 p24 and SARS-Cov-2 S2 (top) and S1 (bottom). **b-c.** quantification of western blots showing c. ratio of spike:p24 in virions, **c**. ratio of S2:total spike**. d.** Western blot of cell lysates used to produce virions with **e**. quantification of S2 to total spike ratio. **f.** brightfield images of lower airway organoids **g**. brightfield images of of cholangiocyte organoids (Scale bars 200 µm).

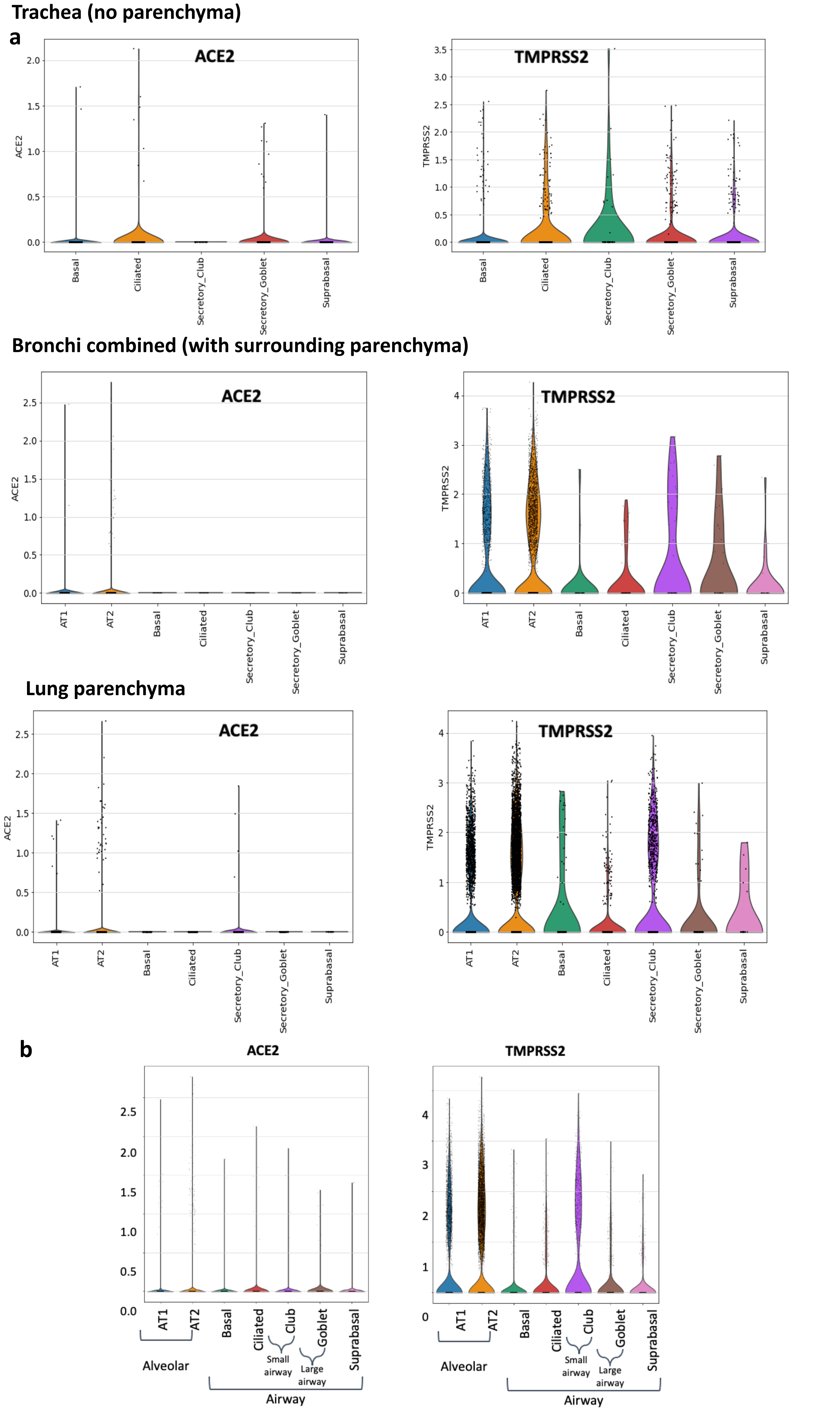


**Supplementary Figure 7: a.** Log-normalised expression of ACE2 and TMPRSS2 genes in single-nuclei RNAseq data for indicated tissue types and. **b.** Log-normalised agreggrated expression of ACE2 and TMPRSS2 genes in single-nuclei RNAseq data from human alveolar (AT1 - alveolar type 1, AT2 - alveolar type 2 pneumocytes), and airway epithelial cells (basal, suprabasal, goblet and ciliated). Data are derived from in Madissoon et al

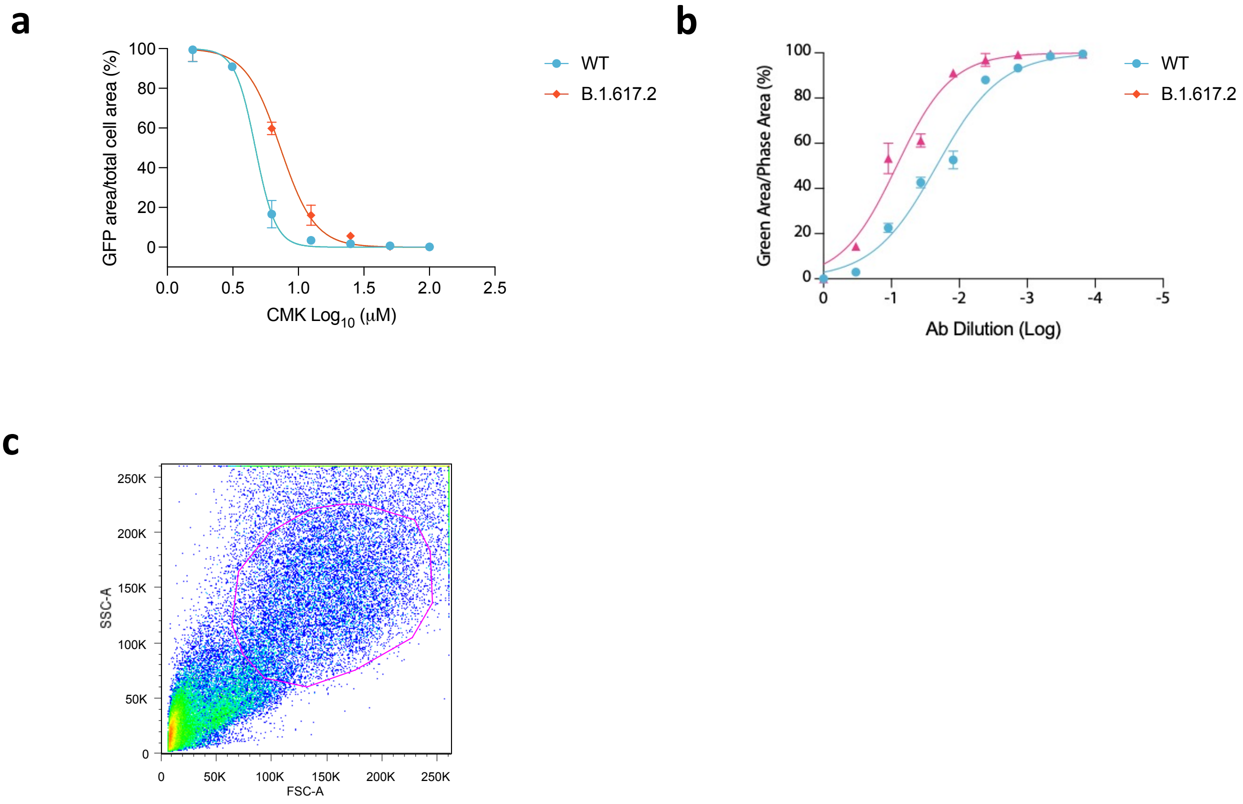


**Supplementary Figure 8: SARS-CoV-2 cell-cell fusion assay and spike expression. a.** effect of furin inhibition using CMK on cell-cell fusion for WT (Wuhan-1 D614G) and Delta variant**.** CMK drug dilution indicated on x axis and fusion on y axis**.** Quantification of cell-cell fusion shows percentage of green area to total cell area. Mean of technical replicates is plotted with error bars representing SEM. Data are representative of at least two independent experiments. **b.** impact of neutralising antibodies from UK wave one 2020 sera on cell-cell fusion for WT (Wuhan-1 D614G) and Delta variant. Serial dilutions of sera added to acceptor cells before co-cultivation with donor cells. **c.** Gating strategy for cell surface staining of SARS-CoV-2 spike proteins from 293T cells transfected with plasmids expressing spike, using rabbit anti-SARS-CoV-2 spike S1/2 polyclonal antibody at 1:100.
